# Single-cell transcriptomics reveals that tumor-infiltrating natural killer cells are activated by localized ablative immunotherapy and share anti-tumor signatures induced by immune checkpoint inhibitors

**DOI:** 10.1101/2023.05.02.539163

**Authors:** Kaili Liu, Negar Sadeghipour, Ashley R. Hoover, Trisha I. Valero, Coline Furrer, Jacob Adams, Abdul Rafeh Naqash, Meng Zhao, James F. Papin, Wei R. Chen

## Abstract

**Rationale:** Natural killer (NK) cells provide protective anti-cancer immunity. However, the cancer therapy induced activation gene signatures and pathways in NK cells remain unclear.

**Methods:** We applied a novel localized ablative immunotherapy (LAIT) by synergizing photothermal therapy (PTT) with intra-tumor delivering of the immunostimulant N-dihydrogalactochitosan (GC), to treat breast cancer using a mammary tumor virus-polyoma middle tumor-antigen (MMTV-PyMT) mouse model. We performed single-cell RNA sequencing (scRNAseq) analysis to unveil the cellular heterogeneity and compare the transcriptional alterations induced by PTT, GC, and LAIT in NK cells within the tumor microenvironment (TME).

**Results:** ScRNAseq showed that NK subtypes, including cycling, activated, interferon-stimulated, and cytotoxic NK cells. Trajectory analysis revealed a route toward activation and cytotoxicity following pseudotime progression. Both GC and LAIT elevated gene expression associated with NK cell activation, cytolytic effectors, activating receptors, IFN pathway components, and cytokines/chemokines in NK subtypes. Single-cell transcriptomics analysis using immune checkpoint inhibitor (ICI)-treated animal and human samples revealed that ICI-induced NK activation and cytotoxicity across several cancer types. Furthermore, ICI-induced NK gene signatures were also induced by LAIT treatment. We also discovered that several types of cancer patients had significantly longer overall survival when they had higher expression of genes in NK cells that were also specifically upregulated by LAIT.

**Conclusion:** Our findings show for the first time that LAIT activates cytotoxicity in NK cells and the upregulated genes positively correlate with beneficial clinical outcomes for cancer patients. More importantly, our results further establish the correlation between the effects of LAIT and ICI on NK cells, hence expanding our understanding of mechanism of LAIT in remodeling TME and shedding light on the potentials of NK cell activation and anti-tumor cytotoxic functions in clinical applications.

## Introduction

Natural killer (NK) cells belong to the innate lymphoid cell family^1, 2^ and are cytotoxic to the virally infected and cancer cells.^3^ NK cells are identified by surface expression of CD56 and lack of CD3. More than 90% of NK cells in circulation are mature NK cells with CD56^dim^ CD16^+^ expression. These cells are highly cytotoxic to the infected or cancerous cells. In contrast, CD56bright CD16^-^ population is found in a much lower percentage in the peripheral blood and mainly resides in lymphoid tissues.^4^ These cells are involved in cytokine production such as IFNγ and TNF.^5^ Secreted IFNγ from NK cells activates nearby immune cells to encourage antigen presentation and adaptive immune responses.^6^ Despite earlier understandings that NK cells react rapidly and non-specifically to malignant cells, recent studies showed that NK cells can differentiate into adaptive cells and generate long-lived immunological memory features.^7^ NK cells express a diverse compilation of activating and inhibitory receptors that conduct their mechanism of action against target cells.^8^ This allows them to have multiple functions to restrict the growth and invasion of cancer cells. In solid tumors, however, tumor-infiltrating NK cells often express low levels of activating receptors and high levels of inhibitory receptors,^9^ consequently, they produce low IFNγ and exert low cytotoxicity.

NK cells have complementary cytotoxic functionality to T cells. Many types of cancer cells have downregulated MHC-I, known as ‘missing-self’, to evade CD8^+^ T cell detection.^10^ Circulating mature NK cells are recruited into the tumor microenvironment by proinflammatory chemokines, and they can identify and efficiently kill tumor cells with low expression of MHC-I. In addition, NK cells are the most important lymphocytes participating in antibody-dependent cell mediated cytotoxicity (ADCC), contributing to cell killing and production of cytokines after administration of therapeutic monoclonal antibodies.^11^ Infiltration and cytotoxicity of NK cells in the cancer tissue influence treatment efficacy and survival,^12^ thus activation of NK cells upon new treatments needs to be evaluated.

A wide variety of in vitro and in vivo platforms have explored the therapeutic capacity of NK cells against cancer, some of which are currently under clinical investigation.^3, 13^ Mature NK cells can be derived from hematopoietic progenitor cells^14^ or human embryonic stem cells^15, 16^ and iPSCs.^17^ The cells can then be proliferated in culture and get prepared in advance, optimized, and activated in the presence of cytokines and stroma for administration into the patients. For example, anti-CD19 chimeric antigen receptor NK cells have enhanced cytotoxic effects against CD19-positive tumors.^18^ CD16 and NKG2C activate adaptive NK cells and increase their proliferation and secretion of IFNγ and enhance ADCC.^19, 20^ Pre-activating of NK cells with IL-12, IL-15, and IL-18 makes them differentiate into memory-like cells.^21^ Moreover, it has been shown that NK cells show activity upon immune checkpoints inhibitor treatments, such as anti- CTLA4 and anti-PDL1.

We have previously developed a localized ablative immunotherapy (LAIT) that combines local photothermal therapy (PTT) and intratumoral injection of a novel immunoadjuvant, N-dilydrogalactochitosan (GC). LAIT extended animal survival in the mouse MMTV-PyMT mammary tumor model^22^ as well as B16-F10 melanoma model.^23^ Using single-cell RNA sequencing (scRNAseq), we showed that LAIT activates sustainable adaptive antitumor responses by increasing the proportions of naïve CD8^+^ T-cells and memory CD4^+^ T-cells from a resting state to an activated state. In addition, DEG analysis revealed that LAIT promoted the expression of genes involved in T cell activation, function, and/or survival, such as Gimap transcripts. Similarly, we observed an increase in the proportion of tumor-infiltrating B cells and differentiation of them from a resting state toward an effector phenotype. Further gene analysis showed that LAIT upregulated genes were involved in B cell activation, BCR signaling, antigen presentation and response. Earlier clustering of scRNAseq from our MMTV-PyMT model revealed subpopulations of NK cells in tumor infiltrated lymphoid cells.^22^ Given the complementary role of NK cells to T-cells and B-cells and their potential as therapeutic immune cells, we were interested in understanding NK cells activation and signaling regulation upon GC, PTT, and LAIT treatment.

In this study, we investigate LAIT’s effect on the antitumor phenotype of NK cells and the correlation between the effects of ICI and LAIT on NK cells in terms of survival of cancer patients, using scRNAseq data form MMTV-PyMT mammary tumor model collected for four different groups: control, GC, PTT, and LAIT. Furthermore, to unleash the full immunological potential of NK cells, we tested the similarities between genes activated after LAIT treatment from our study and various cancer patients after ICI treatment. Our study for the first time showed that LAIT activates cytotoxicity in NK cells and the upregulated genes positively correlate with beneficial clinical outcomes for cancer patients. More importantly, our results further establish the correlation between the effects of LAIT and ICI on NK cells, hence expanding our understanding of mechanism of LAIT in remodeling TME and shedding light on the potentials of NK cell activation and anti-tumor cytotoxic functions in clinical applications.

## Results

### ScRNA-seq reveals heterogeneity of tumor-infiltrating natural killer cells upon LAIT

We previously reported the promising therapeutic effects of localized ablative immunotherapy (LAIT) on MMTV-PyMT breast tumors and uncovered the anti-tumor roles of tumor-infiltrating B cells^24^ and T cells^25^ in response to LAIT. Although there are many other reports revealing the transcriptional atlas and anti-tumor roles of both innate and adaptive immune cells by scRNAseq, the contributions of NK cells were less known. To address this, we *in silico* subset the NK cells by choosing CD3^-^NK1.1^+^ (*Cd3g^-^/d^-^/e^-^Klrb1c^+^*) or CD3^-^NKp46^+^ (*Cd3g^-^/d^-^/e^-^Ncr1*^+^) cells from tumor-infiltrating lymphoid cell compartments^25^ and obtained 1,257 NK cells, using scRNAseq (Figure S1A). Feature plot showed that these cells were abundantly expressed with NK markers *Klrb1c* and *Ncr1* (Figure S1B). Although they had sparse/sporadic expressions of *Cd8a/b1* or *Cd4*, they had no expression of T cell markers *Cd3g/d/e*, proving that they were NK cells without mixture of T cells. Single-cell re-clustering resulted in 5 clusters (C0-C4) (Figure 1A), among which the cell number and proportion in each treatment group were calculated (Figure 1B-C). We obtained cluster-specific gene markers (Figure S1C) and found cells in C2 (Figure 1A) were enriched with *Gzmb* and *Prf1* and therefore annotated as cytotoxic NK cell subtype (Cyto.NK). C3 was cycling NK (Cyc.NK) due to specific expression of proliferating marker *Mki67* and *Top2a*. C4 was enriched with interferon signaling genes (*Ifit3*, *Ifit1*, *Ifi209*, *Isg15*) and was classified as interferon-stimulated NK (IFN-NK) with inflamed properties. We named C0 and C1 as activated NK cells (Act.NK) because activating NK receptor *Klra4* was expressed in C0 (Figure S1C) and both clusters were expressed with *Tnf* and *Ifng* (Figure 1D).

**Figure 1.**
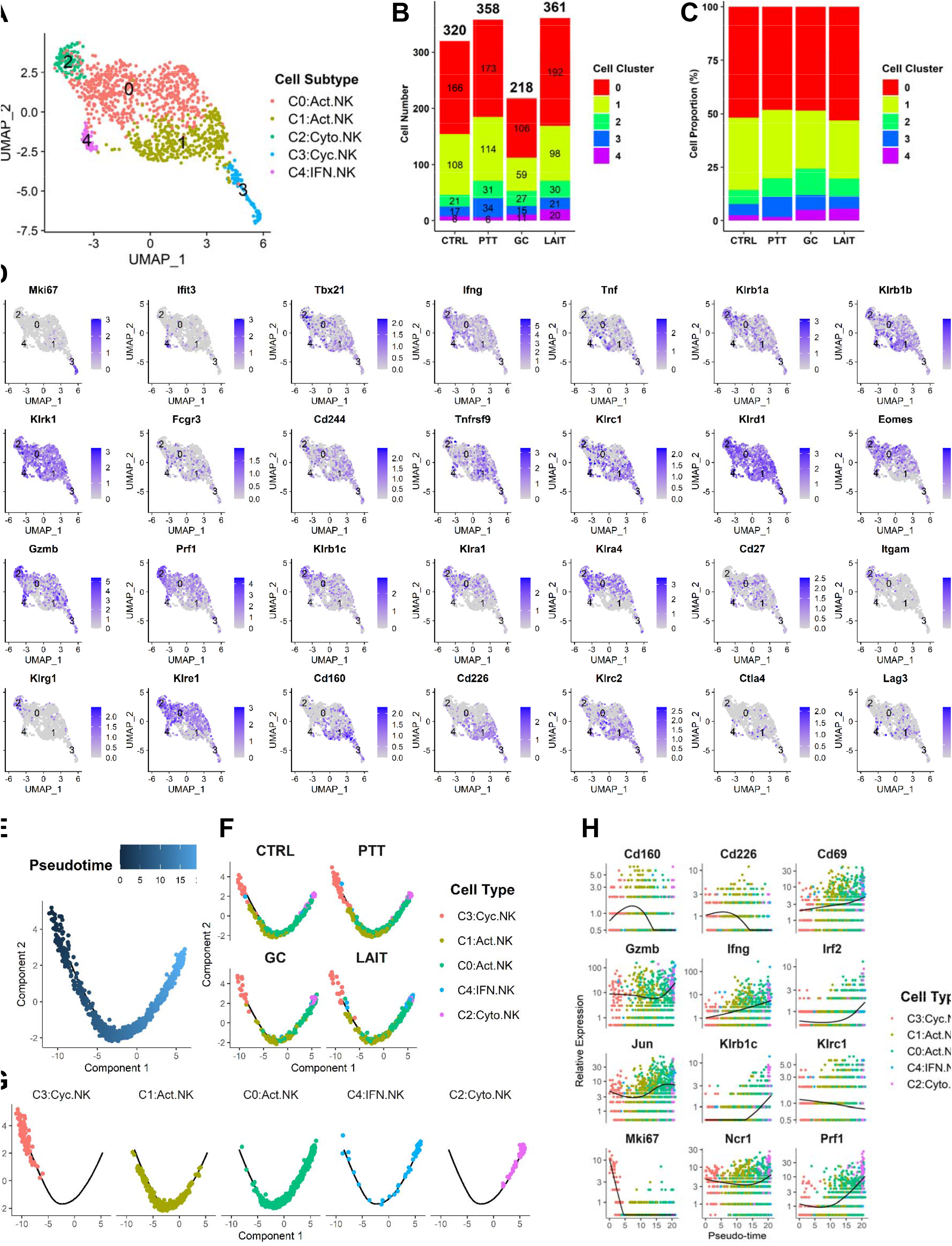
Analysis of tumor-infiltrating natural killer cells in different treatment groups (Control, PTT, GC, and LAIT). **(A)** Two-dimensional visualization of single-cell clusters using method of Uniform Manifold Approximation and Projection (UMAP) for tumor-infiltrating natural killer (TINK) cells from all treatment groups. TINK cells were classified into cycling NK (C3), activated NK (C1, C2), Inflamed/IFN-NK (C4), cytotoxic NK (C2). **(B)** Bar plot showing the NK cell number in each cluster in each treatment. The total cell number for each treatment was labelled on top of bar. **(C)** Bar plot showing the relative NK cell proportion in each cluster in different treatment groups (CTRL, PTT, GC, and LAIT). The proportion was calculated by using NK cell number in each cluster divided by the total number within given treatment. **(D)** Feature plots showing expressions of selected NK cell markers. **(E)** Cell trajectory plot of NK cells colored by pseudotime. **(F)** Cell trajectory plots of NK cells colored by subtypes in each treatment group. **(G)** Cell trajectory plots of each NK cell subtype from all treatment groups. **(H)** Selected gene expression plots following pseudotime colored by NK cell subtype.

To validate above cell subtype annotations, we utilized a panel of NK cell markers and examined their expression patterns in NK cell clusters (Figure 1D). Transcription factor T-bet (*Tbx21*), cytokine genes *Ifng* and *Tnf* were ubiquitously expressed in all NK cell clusters. This broad gene expression pattern was also found in activating NK cell receptor genes *Klrb1a/b*, NKG2D/CD314 (*Klrk1*), CD16 (*Fcgr3*), 2B4/SLAM4 (*Cd244*), CD137/4-1BB (*Tnfrsf9*), and inhibitory NK cell receptors NKG2A (*Klrc1*) and CD94 (*Klrd1*) (Figure 1D). *Eomes*, *Gzmb*, *Prf1*, activating receptors *Klrb1c* and *Klre1*, inhibitory receptors *Klra1/4* were rarely expressed in C1 while highly expressed in other remaining clusters (C0, C2-4), suggesting that in both activated NK cell clusters of C0 and C1, C0 may be more activated than C1. CD11b (*Itgam*) was enriched in C2 and showed a complementary expression pattern to *Cd27*. Since it has been reported that CD27^bright^ NK cells had greater cytotoxic potential and produced higher levels of cytokines than CD27^dim^ NK cells and CD27^-^CD11b^+^ (*Cd27^-^Itgam^+^*) type represents mature NK state while CD27^+^CD11b^-^ (*Cd27^+^Itgam^-^*) marks immature type of NK, it demonstrated that C2 was mature while other clusters were immature/less mature. This agreed with our annotation that C2 was cytotoxic NK with mature effector functions while other clusters were immature with proliferating and activation states. We also found that expressions of *Cd160*, *Cd226*, inhibitory receptor *Klrc2*, immune checkpoint inhibitor *Ctla4* and *Lag3* were lower in C2 while higher in other clusters. Expressions of NK inhibitory receptor CD158 (*Kir3dl1/2*) or checkpoint inhibitor PD1 (*Pdcd1*) or TIM3 (*Havcr2*) were not detected.

To further confirm the scRNAseq clustering and NK cell subtype annotation, we performed single-cell trajectory inference^26^ and found NK cell subtypes were localized with distinct and overlapping patterns along the trajectory (Figure 1E-G). Following the pseudotime progression (Figure 1E), cycling NKs (C3) were localized at the start of trajectory, with activated NKs (C1, C0) as well as IFN-NKs (C4) in the middle trajectory, and cytotoxic NKs (C2) distributed at the end (Figure 1G). Gene expressions along pseudotime revealed that *Mki67* and inhibitory receptor NKG2A (*Klrc1*) were expressed highest in the early pseudotime where cycling NK cells reside and showed a continuous decreasing expression trend till end of pseudotime (C2). NK activating receptors *Cd160* and DNAM1 (*Cd226*) were highly expressed in the middle of this trajectory where activated NK cells (C1 and C0) were localized (Figure 1H). NK activating receptors *Klrb1c* and *Ncr1* displayed unique and high expressions in C4 and C2, in a pattern the same as cytotoxic markers *Gzmb* and *Prf1* (Figure 1H). Activation markers *Cd69* and anti-tumor cytokine *Ifng* showed a continuously rising expression pattern ranging from cycling, activated to cytotoxic NKs, suggesting that the pseudotime proceeds toward the activation and effector direction (Figure 1H). The coupling of pseudotime with NK activation/cytotoxicity process were further supported by expressions of more NK activating gene signatures in the heatmap (Figure S1F).

Next, we investigated the constitution and distribution of NK subtypes in each treatment group (Figure S1D). By calculating the NK cell proportions from the total CD45^+^ immune cells, we found LAIT increased the proportions of interferon-stimulated NK cells (C4, 0.18%) when compared with control (0.07%), although no significant induction in remaining NK subtypes by treatment of PTT, GC, or LAIT (Figure 1C, S1E) was observed. This result also prompted us to investigate the potential NK cell’s transcriptional alterations by LAIT.

### Differential gene expression and functional enrichment analysis of tumor-infiltrating NK cells

scRNAseq can not only reveal the cell composition heterogeneity but also harvest the transcriptome of individual cells, broadening the horizon of research in physiology, pathology and therapy^27–31^. To explore the effect of each treatment on the functions of tumor-infiltrating NK cells at the transcriptional level, we performed differential gene expression and functional enrichment analysis. Three comparisons were adopted: PTT vs CTRL, GC vs CTRL, and LAIT vs CTRL. For each comparison, differentially expressed genes (DEGs) were generated by using *FindMarkers* function in *Seurat* R package^32^. DEGs from comparing two treatment groups were defined as log Fold change > 0.25 for the upregulated and < –0.25 for the downregulated along with adjusted *p* value < 0.05, shown as red and blue dots in the volcano plots, respectively (Figure 2A-C). Functional enrichment analyses were carried out using both over-representation analysis (ORA)^33^ and gene set enrichment analysis (GSEA)^34^. The *clusterProfiler* R package^35^ was chosen to conduct and visualize functional enrichment analysis by using databases of Gene Ontology (GO)^36^, Kyoto Encyclopedia of Genes and Genomes (KEGG)^37^, Reactome Pathway Database (Reactome)^38^ and Molecular Signatures Database (MsigDB)^39^.

**Figure 2.**
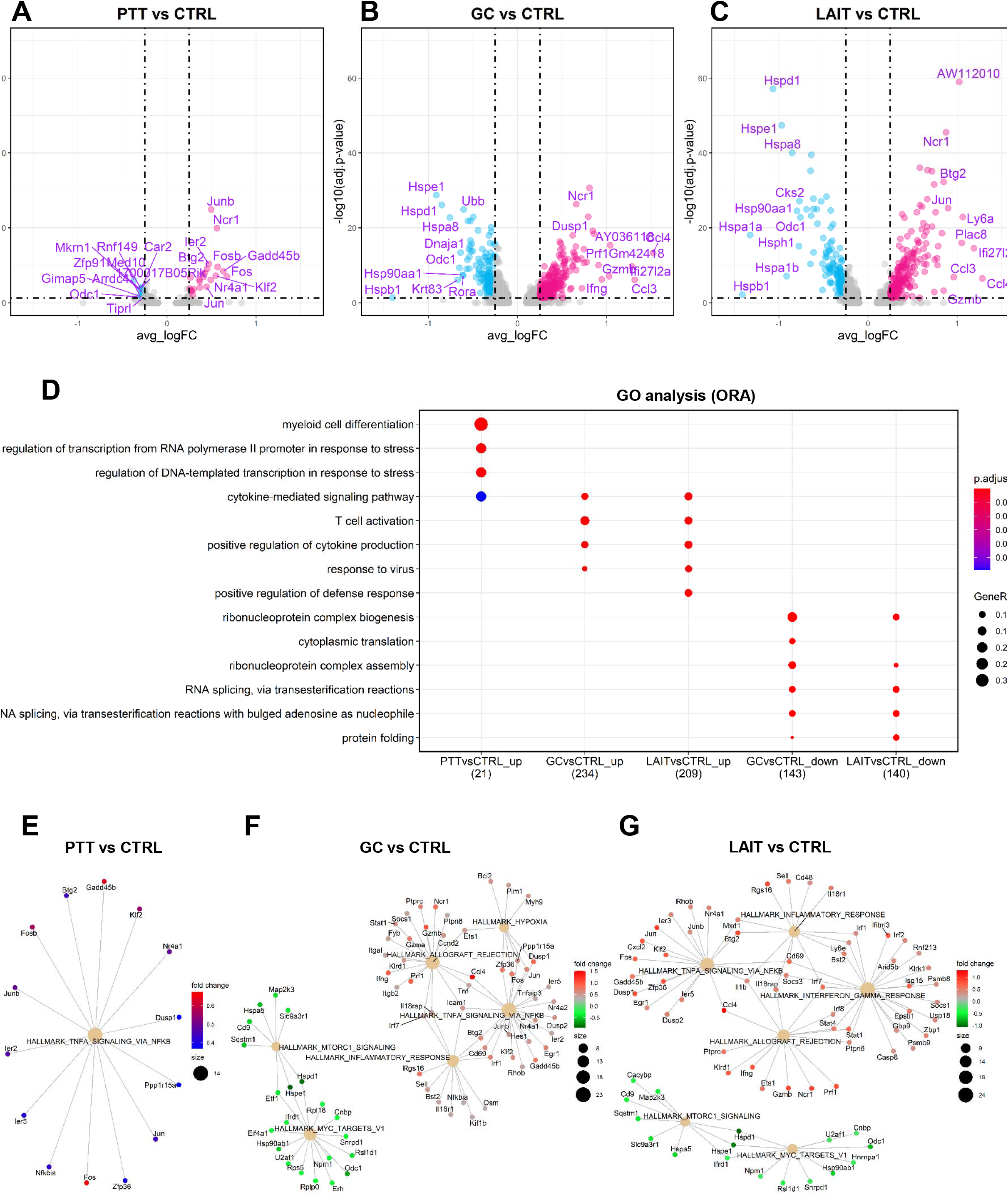
Differential gene expression and pathway enrichment analysis for TINK cells. **(A)** Volcano plot showing differential gene expression comparing PTT with CTRL (PTT vs CTRL). Top 10 upregulated and downregulated genes are labeled. **(B)** Volcano plot showing differential gene expression comparing GC with CTRL. **(C)** Volcano plot showing differential gene expression comparing LAIT with CTRL. **(D)** Dot plot showing biological process (BP) of gene ontology (GO) analysis using over-representation analysis (ORA) method for both upregulated (up) and downregulated (down) genes for PTT vs CTRL, GC vs CTRL, and LAIT vs CTRL. **(E)** Network plot showing gene set enrichment analysis (GSEA) for DEGs from PTT vs CTRL using MsigDB hallmark gene sets. **(F)** Network plot showing the GSEA for DEGs from GC vs CTRL. **(G)** Network plot showing the GSEA for DEGs from LAIT vs CTRL.

For PTT vs CTRL, 22 upregulated and 15 downregulated genes were shown in the volcano plot (Figure 2A). Biological process (BP) of gene ontology (GO) analysis using the ORA method demonstrated that PTT enriched myeloid cell differentiation, transcription in response to stress and cytokine-mediated signaling pathways compared to CTRL (Figure 2D). This result was supported by KEGG analyses with TNF signaling and NK cell mediated cytotoxicity process (Figure S2A) and Reactome analyses with TLR3/10 cascade and interleukins signaling (Figure S2B). In contrast, GO enrichment of PTT downregulated genes was not found (Figure 2D). KEGG and Reactome enrichment analyses showed that PTT downregulated genes were enriched in metabolism related signaling pathways (Figure S2A-B).

GC treatment upregulated 246 genes and downregulated 151 genes compared to CTRL (Figure 2B). GO analysis revealed that GC upregulated genes were enriched in cytokine-mediated signaling, T cell activation, positive regulation of cytokine production, and response to virus (Figure 2D). Similar results were obtained by using KEGG and Reactome analyses and displayed overlap with PTT-induced pathways (Figure S2A-B). Conversely, GC downregulated genes were involved in ribosome, spliceosome, nonsense-mediated decay pathways (Figure 2D, Figure S2A-B).

When comparing LAIT vs CTRL, we observed 220 upregulated and 151 downregulated genes (Figure 2C). LAIT-upregulated genes were enriched in positive regulation of defense response and many processes that were observed by GC treatment (Figure 2D, Figure S2A-B). Enrichment of LAIT downregulated genes were like those in GC, also involved in ribonucleoprotein complex biogenesis/assembly, RNA splicing, protein folding/processing (Figure 2D, Figure S2A), spliceosome, and chaperon cycle pathways (Figure S2B).

The ORA analysis revealed that PTT, GC and LAIT owned overlapping gene expression signatures (Figure 2D), although the numbers and fold changes of DEGs were not identical (Figure 2A-C). To further explore gene expression differences induced by these treatments, we utilized hallmark gene sets of MsigDB and carried out GSEA to visualize gene-pathway networks (Figure 2E-G). As shown in Figure 2E, PTT elevated TNFα signaling. GC enriched hypoxia, allograft rejection, TNFα signaling and inflammatory responses while downregulated Myc targets and mTORC1 pathways (Figure 2F). Like GC, LAIT upregulated TNFα signaling, allograft rejection, and TNFα signaling. Moreover, it specifically enriched IFNγ signaling, indicating that the anti-tumor cytotoxicity and effector functions were triggered by LAIT in NK cells. LAIT shared the downregulated Myc targets and mTORC1 signaling with GC (Figure 2G). These analyses showed that PTT, GC and LAIT had overlapping pathway such as TNFα response. However, GC and LAIT enriched more proinflammatory pathways and had more associated genes than those enriched by PTT alone.

In addition to the MsigDb collection, the KEGG and Reactome pathway databases were chosen for GSEA. For PTT vs CTRL, no gene set enrichment was found, most likely due to the small number of DEGs between the two groups. GC treatment upregulated pathways involved NK cell mediated cytotoxicity, leishmania infection, and T cell/Toll-like receptor signaling pathways (Figure S2C) and IFNγ signaling (Figure S2E) and downregulated ribosome, spliceosome (Figure S2C), autophagy, and other stress related metabolism pathways including cellular response to starvation, amino acid deficiency and protein translation elongation pathways (Figure S2E). LAIT upregulated signaling pathways of NK cell mediated cytotoxicity, cytokine/chemokine (Figure S2D), IFNαβ/γ (Figure S2F), and downregulated MAPK, spliceosome (Figure S2D) and heat shock protein composed pathways (Figure S2F). Taken together, both ORA and GSEA led to consistent enrichment results (Figure 2D-G, S2A-F). It also demonstrated that treatments of PTT, GC, and LAIT had overlapping and nonoverlapping functions on NK cells. This was further elucidated by the degrees of NK cell activation from individual treatments to the combined LAIT treatment (Figure 2E-G, S2C-F). These results provided evidence that LAIT had a greater influence on activating NK cell gene signatures and potential anti-tumor functions than PTT or GC alone.

### Overlapping of treatment-upregulated genes in tumor-infiltrating NK cells

We hypothesized that understanding the relationships among the treatment regulated DEGs would shed light on the molecular mechanism stimulated by LAIT therapy. To determine NK cell gene expression signatures associated with LAIT-enhanced survival of tumor-bearing mice, we investigated the relationship of treatment-regulated DEGs and focused on the LAIT-specific DEGs in NK cells. A Venn diagram was used to display the overlap of PTT, GC, LAIT regulated genes (Figure 3A). Genes uniquely regulated by PTT alone were confined in Set_0. PTT-GC- (LAIT) intersecting genes were confined in Set_1; GC-(LAIT) intersecting genes (excluding Set_1) were confined in Set_2. Genes uniquely regulated by GC alone were grouped in Set_3. Similarly, Set_0, Set_3 and Set_4 was defined as specific upregulation by PTT, GC and LAIT, respectively.

**Figure 3.**
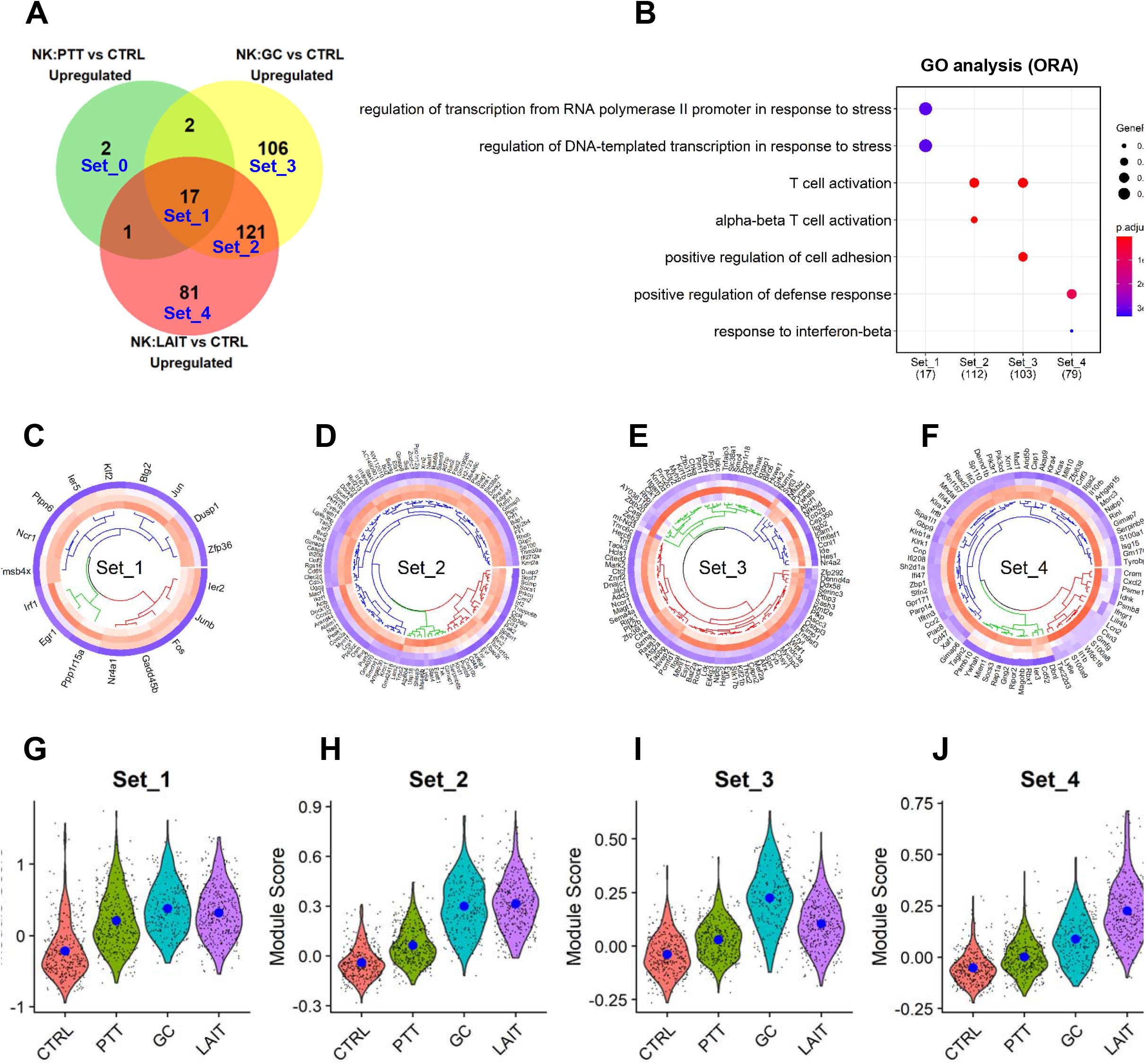
Overlapping of treatment-upregulated genes in TINK cells. **(A)** Venn diagram showing upregulated genes from comparisons of PTT vs CTRL, GC vs CTRL, and LAIT vs CTRL. Five gene sets, from Set_0 to Set_4, are labeled. **(B)** Dot plot for BP of GO analysis of genes in Set_1 to Set_4. **(C)** Circular heatmap showing the expression of genes from upregulated Set_1 in each treatment group. Heatmap columns for groups of CTRL, PTT, GC and LAIT were arranged from outside to inside. Higher expression was colored in red while lower in blue. **(D)** Circular heatmap showing the expression of genes from upregulated Set_2 in each treatment group. **(E)** Circular heatmap showing the expression of genes from upregulated Set_3 in each treatment group. **(F)** Circular heatmap showing the expression of genes from upregulated Set_4 in each treatment group. **(G)** Violin plot showing relative gene signature module scores from Set_1 in each treatment group. **(H)** Violin plot showing relative gene signature module scores from Set_2 in each treatment group. **(I)** Violin plot showing relative gene signature module scores from Set_3 in each treatment group. **(J)** Violin plot showing relative gene signature module scores from Set_4 in each treatment group.

Collectively, 17 genes of Set_1 were upregulated among PTT, GC, and LAIT (Figure 3A). Individually, PTT, GC and LAIT specifically upregulated 2 (Set_0), 106 (Set_3) and 81 (Set_4) genes, respectively (Figure 3A). GO analysis for the individual gene sets were shown in Figure 3B. Biological processes (BP), such as transcription in response to stress response, was enriched in Set_1 which were common to all 3 treatments (Figure 3B). (αβ)T cell activation was enriched in Set_2, only shared by GC and LAIT (Figure 3B). GC alone (Set_3) enriched pathways of T cell activation and positive regulation of cell adhesion (Figure 3B). LAIT-specific (Set_4) pathways were enriched for positive regulation of the defense response and response to IFNβ (Figure 3B).

To confirm the above enrichment analysis for different gene sets, MsigDB, KEGG, and Reactome analyses were used. Similar enrichments were found (Figure S3A-C) compared with GO analysis (Figure 3B). For example, LAIT-specific genes (Set_4) were involved with IFNγ/α responses (Figure S3A), NK cell mediated cytotoxicity (Figure S3B).

Although enrichments from genes in Set_1 to Set_4 share NK cell activation related pathways, we hypothesized that their capacities for activating NK cells were different for each treatment. To test this hypothesis, we compared the expression levels of each component of these confined gene sets in different treatment groups using circular heatmap. Set_0 was not used for analysis likely due to the small number of genes. Even though Set_1 genes were common to all treatment groups, levels of most genes in groups of GC and LAIT were higher than those in CTRL and PTT (Figure 3C). This was also observed in GC and LAIT shared gene Set_2 (Figure 3D). Consistent with the pattern established in Set_1 and Set_2, Set_4 genes showed the highest expression in NK cells in the LAIT treated tumors (Figure 3F). In GC-specific upregulated gene Set_3, GC treatment showed the highest stimulation (Figure 3E).

Instead of showing the individual expression of genes from Set_1 to Set_4 in each treatment, we next used *AddModuleScore* function from *Seurat* R package^32^ to calculate the average expression level of these upregulated gene sets. These module scores for each treatment were displayed using violin plots with each dot representing a score of a single cell (Figure 3G-J). The expression scores in gene Set_1 (Figure 3G) and Set_2 (Figure 3H) for GC and LAIT treatments were higher than that in CTRL and PTT. GC has the highest module score in Set_3 (Figure 3I) because this gene set is GC-specifically upregulated. Consistently, LAIT has the highest NK cell activation degree in Set_4 because this gene set is specifically upregulated by LAIT (Figure 3J), also indicating that PTT and GC can synergize to stimulate the expression of gene signatures in Set_4. This result is consistent with that of the gene expression heatmaps (Figure 3C-F) and consistent with the hypothesis that although genes in Set_1, Set_2, and Set_4 were all enriched in NK cell activation related pathways, LAIT resulted in higher degrees of NK cell activation than GC or PTT alone.

Next, we utilized the published gene signatures of pro-inflammatory function, cytolytic effector, Type I/II IFNs that have been reported in human breast cancer^40^ and detected their module expression level in response to LAIT therapy in our breast tumor model. We found that GC drove the module score of these signaling pathways and made the GC and LAIT groups have highest module scores (Figure 3K-N). This suggests that LAIT enables upregulation of type I/II IFN and pro-inflammatory pathways and contribute to the induction of cytotoxic tumor killing effect in NK cells.

To further investigate the effect of LAIT on transcriptional changes in NK subpopulations, we selected a panel of representative genes related to NK functions and compared their expressions in each treatment group across NK subtypes. These genes were NK activation/differentiation, cytolytic effector molecules, activating receptors, inhibitory receptors, IFN response pathway components, cytokine/chemokines (Figure S2D). Activation related markers *Cd69* and *Cd44* were upregulated in early cycling and activated NK cells with an increasing trend by PTT, GC and LAIT. NK differentiation marker *Tbx21* and *Sell* were induced by treatments in activated and more matured IFN and cytotoxic NKs. Stress response markers *Fos* and *Jun* were induced by treatments in all NK subtypes. Cytotoxic markers *Gzma* and *Gzmb* were sharply induced by treatments in IFN NKs and reached a high level in cytotoxic NK cells, while *Gzmc* was mainly expressed and induced by LAIT in cycling and early activated NK cells (C0). Activating receptors of *Klra4*, *Klrb1a/c*, *Klre1*, *Klrk1*, *Ncr1*, and *Slamf7* were mainly expressed and upregulated by GC and LAIT in cytotoxic NK subtypes, although many were also induced by LAIT in in activated, and IFN-NK cells. *Cd27* and *Tnfrsf9* were induced by treatments while mainly expressed non-cytotoxic NK subpopulations. Inhibitory receptor/immune checkpoint *Ctla4* and *Lag3* were both induced by LAIT in non-cytotoxic NKs. However, in cytotoxic NKs, *Ctla4* was induced while *Lag3* was inhibited by LAIT. Interferon response pathway genes, including *Ddx58/60*, *Stat1/2/4*, *Irf1/2/7*, *Ifi203/206/208/209* were mainly expressed in IFN-NK cells and driven by GC. *Ifi27l2a* and *Ifitm3* were not only expressed and induced by LAIT in IFN-NK, but also in cycling and activated NK cells. *Ccl3/4/5* were mainly expressed and induced by GC and LAIT in IFN- and cytotoxic NKs. *Ccr2* was induced by LAIT in all NK subtypes while *Cxcl10* was induced by LAIT mainly in cycling and IFN-NK cells. This analysis suggested that LAIT triggered the expressions of NK activation gene signatures, activating receptors and pro-inflammatory cytokines/chemokines and IFN pathways, this is consistent with the module score results (Figure 5A).

### Overlapping of treatment-downregulated genes in tumor-infiltrating NK cells

We next investigated the relationship of treatment-downregulated DEGs and focused on the LAIT-specific negative regulation on these genes in tumor-infiltrating NK cells. A Venn diagram was used to illustrate the overlap of PTT, GC, and LAIT downregulated genes (Figure 4A). Genes intersected with different treatments were grouped similarly to Figure 3A: Set_0 was for PTT-specific genes; Set_1 for PTT-GC-(LAIT) intersecting genes; Set_2 for GC-(LAIT) intersected genes (Set_1 excluded), Set_3 for GC-specific genes, and Set_4 for LAIT-specific genes. (Figure 4A).

**Figure 4.**
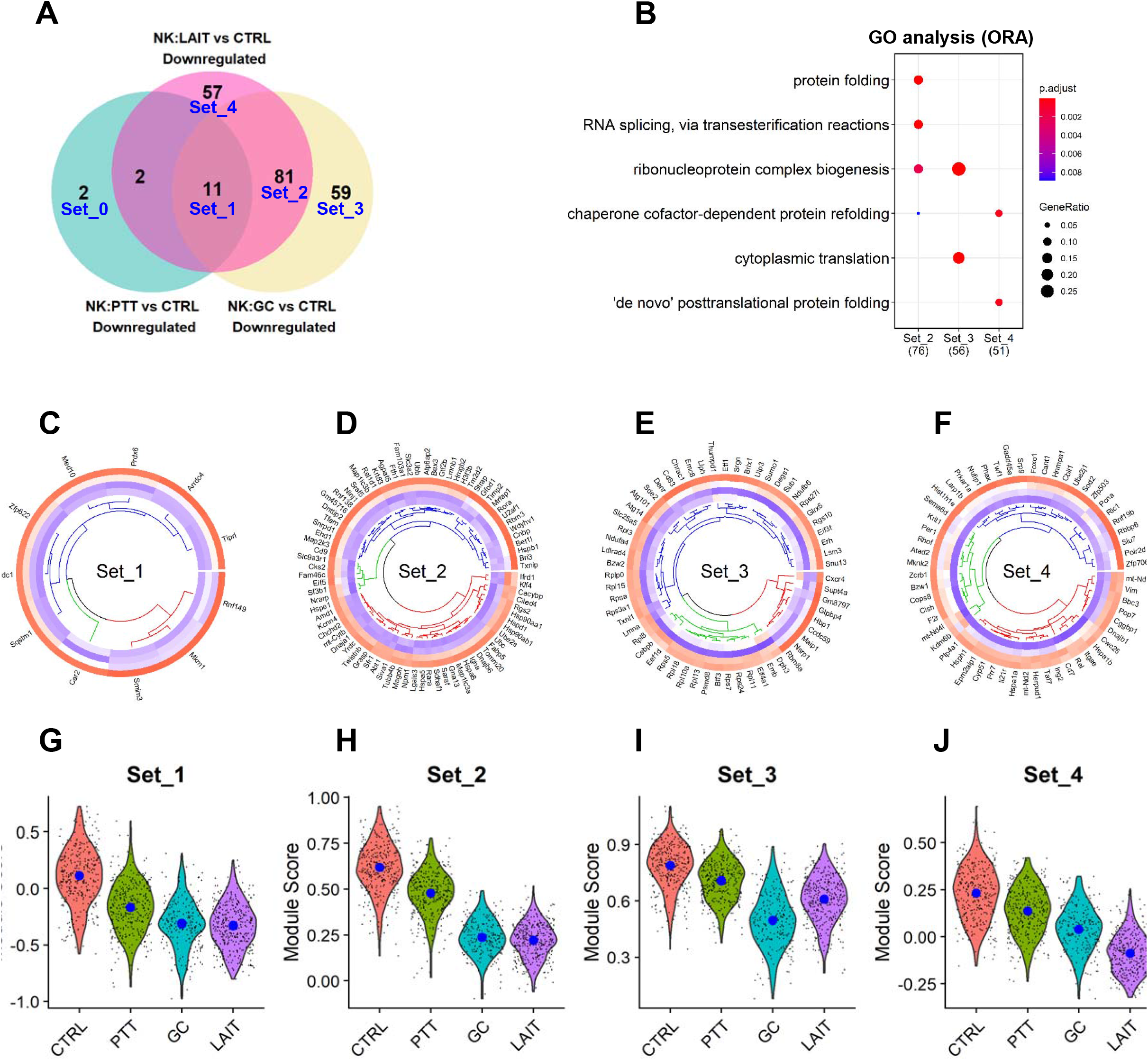
Overlapping of treatment-downregulated genes in TINK cells. **(A)** Venn diagram showing downregulated genes from comparisons of PTT vs CTRL, GC vs CTRL and LAIT vs CTRL. Five gene sets, from Set_0 to Set_4, were labeled. **(B)** Dot plot for BP of GO analysis of genes in Set_1 to Set_4. **(C)** Circular heatmap showing the expression of genes from downregulated Set_1 in each treatment group. Heatmap columns for groups of CTRL, PTT, GC and LAIT were arranged from outside to inside. Higher expression was colored in red while lower in blue. **(D)** Circular heatmap showing the expression of genes from downregulated Set_2 in each treatment group. **(E)** Circular heatmap showing the expression of genes from downregulated Set_3 in each treatment group. **(F)** Circular heatmap showing the expression of genes from downregulated Set_4 in each treatment group. **(G)** Violin plot showing relative gene signature module scores from Set_1 in each treatment group. **(H)** Violin plot showing relative gene signature module scores from Set_2 in each treatment group. **(I)** Violin plot showing relative gene signature module scores from Set_3 in each treatment group. **(J)** Violin plot showing relative gene signature module scores from Set_4 in each treatment group.

Collectively, for Set_1, PTT, GC and LAIT commonly downregulated 11 genes (Figure 4A). GC and LAIT (Set_2) co-downregulated 81 genes, while uniquely, PTT downregulated 2 genes (Set_0), GC downregulated 59 genes (Set_3), and LAIT downregulated 57 genes (Figure 4A). GO analysis of genes from the different sets was shown in Figure 4B. Protein folding/refolding, RNA splicing, and ribonucleoprotein complex biogenesis pathways were enriched in Set_2. Set_3 and Set_4 were enriched in pathways involved in ribonucleoprotein complex biogenesis and protein translation/folding (Figure 4B). The results in Figure 4A-B demonstrate that PTT, GC and LAIT treatments shared some overlapping gene regulation, like what was observed for the upregulated genes in Figure 3. Using MsigDB (Figure S4A), KEGG (Figure S4B) and Reactome (Figure S4C) databases disclosed similar findings.

Although protein folding and RNA splicing related pathways were shared among above gene sets, we assumed that their capacities for negatively regulating activated NK cells were distinct within and unique to each treatment. To test this assumption, we compared the expression levels of these confined gene sets in different treatment groups using circular heatmap analysis. Set_0 was not used for analysis likely due to the small number of genes. Gene expression pattern showed an increasing inhibition from CTRL to PTT, GC and LAIT in Set_1, Set_2 and Set_4, suggesting the unique and combinatory effect of each treatment in regulating these gene expression profiles (Figure 4C, 4D, 4F). In Set_3, GC showed the highest inhibition due to these genes were GC-specific downregulated genes (Figure 4E).

Next, we calculated the average expression level of these downregulated gene modules from Set_1 to Set_4 in each treatment (Figure 4G-J). The expression scores for gene Set_1 (Figure 4G) and Set_2 (Figure 4H) were lower in GC and LAIT than that in CTRL and PTT, indicating that the degree of protein folding and RNA splicing from Set_1 and Set_2 genes were predominantly inhibited by GC and LAIT. GC showed the lowest downregulation on expression score in Set_3 (Figure 4I) while LAIT exhibited the lowest module score in Set_4 (Figure 4J). We also utilized the reported anti-inflammatory gene signatures in human breast cancer and tested the expressions in our model in each treatment group. We found treatment of PTT, GC, and LAIT all downregulated the anti-inflammatory gene signatures, therefore contributing to a pro-inflammatory state beneficial for NK cell’s tumor-killing capacity (Figure 5A).

**Figure 5.**
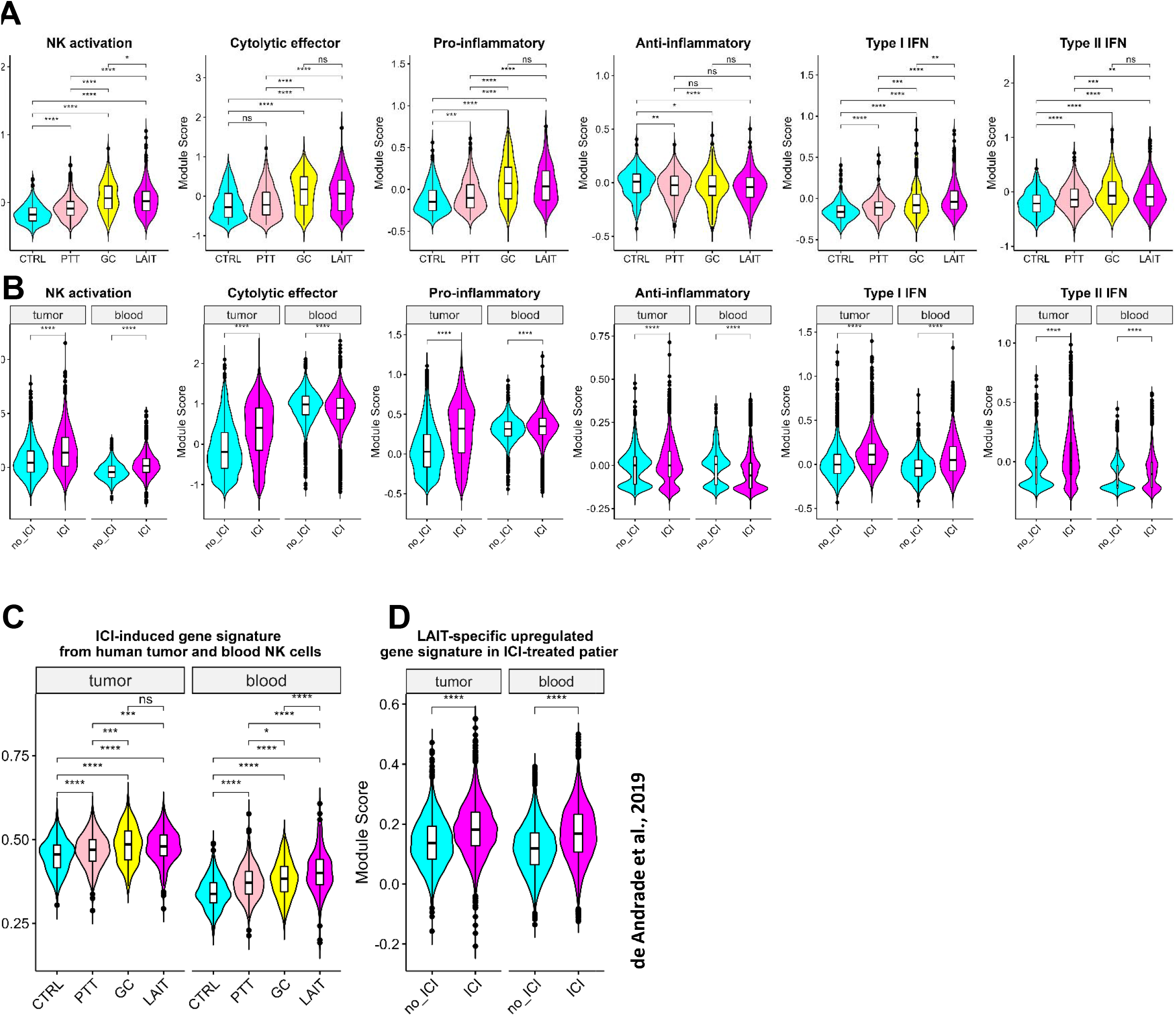
ScRNAseq analysis for similarity between LAIT and immune checkpoint inhibitor (ICI) in stimulating anti-tumor phenotype of NK cells. **(A)** Violin plot showing module scores for anti-tumor phenotype related gene signatures including NK activation, cytolytic effector, pro-inflammatory, anti-inflammatory, type I interferon, and type II interferon gene signatures in treatment groups of CTRL, PTT, GC, and LAIT in mouse PyMT TINK cells. **(B)** Violin plot showing module scores for anti-tumor phenotype related gene signatures including NK activation, cytolytic effector, pro-inflammatory, anti-inflammatory, type I interferon, and type II interferon gene signatures in ICI treated TINK cells and blood NK cells from breast cancer patients. no_ICI represents patient samples without ICI treatment. **(C)** Violin plot showing module scores for ICI-induced gene signatures mapped into treatment groups of CTRL, PTT, GC, and LAIT in mouse PyMT TINK cells. ICI-induced gene signatures from breast cancer patient TINK cells and blood NK cells also demonstrated the increased production in GC/LAIT group, indicating the similar effect of ICI and LAIT. **(D)** Violin plot showing module scores for LAIT specifically-induced gene signatures (upregulated Set_4) mapped into human breast cancer TINK cells and blood NK cells upon ICI treatment. LAIT specifically-induced gene signatures from PyMT mouse TINK cells demonstrated the increased production in ICI treatment group, further confirming the similar effect of ICI and LAIT in stimulating NK cell’s anti-tumor immunity.

Next, we investigated the synergizing effect between PTT and GC in downregulating gene expressions in NK cell subtypes (Figure S4D). We found that NK marker *Cd7* was downregulated by LAIT in non-cytotoxic NK cells and CD16 (*Fcgr3*) downregulated by LAIT in activated, inflamed and cytotoxic NK cells. Immune suppressing factors *Foxo1* and *Tgfb1* were inhibited by LAIT, especially in mature NK subpopulations. Although *Hif1a* was upregulated in cycling and activated NKs, it was downregulated in cytotoxic NKs by LAIT. Heat shock protein transcripts were downregulated mainly by GC and LAIT.

### LAIT-upregulated gene signatures in TINK cells resemble immune checkpoint inhibitor (ICI) treatment in several tumor types

Immune checkpoint inhibitor (ICI) therapy achieved huge progress in tumor immunology. However, whether and how they affect the NK cells are not clear. To address this, we performed the data mining on scRNAseq from ICI treated human patients and mice with multiple cancer types.^41–44^ We found in ICI-treated, patients with melanoma (Figure 5B), breast cancer (Figure S5B) and LMD patients (Figure S5C), and mouse breast cancer model (Figure S5A), gene signatures of NK activation, cytolytic effector, pro-inflammatory, type I and II interferon were increased in ICI group when compared with no ICI treated controls. Meanwhile, anti-inflammatory gene signatures were downregulated. These results suggested that ICI was able to induce NK cell antitumor activation and cytotoxicity, resembling LAIT. To provide a more direct relationship between LAIT and ICI in affecting NK cell gene expressions, we calculated the LAIT-induced DEGs and detected their expressions in our LAIT-treated mice NK cells, we found treatments, especially LAIT/GC showed highest in inducing their expressions (Figure 5C, S5D, F, H). This analysis established a positive correlation between the LAIT and ICI in activating the NK cells at the single cell-based transcriptomic level. In parallel, we mapped the LAIT-specifically upregulated gene set 4 and examined their expressions in ICI treated human cancer patient NK cells. We found expressions of LAIT-specific gene set 4 were also induced by ICI (Figure 5D, S5E,G,I). This result further confirmed our finding that LAIT and ICI shared similarities in modulating NK cell gene expression in TME, suggesting that the combinations of LAIT and ICI has the potential to provide more capacity in exerting anti-tumor functions through NK cells.

### LAIT-upregulated genes in TINK cells positively correlate with greater overall survival in cancer patients

Above results indicated that LAIT drove the activation, IFN response and cytotoxic pathway gene signatures in TINK cells. A combination of PTT with GC augmented these antitumor responses than either PTT or GC alone and achieved better therapeutic efficacy. To study the clinical relevance of the outcome of LAIT’s modulation on TINK cells, we tested the potential relations between LAIT induced gene expressions and cancer patient survival. First of all, we used LAIT specific upregulated and downregulated genes from mouse NK cells, mapped them into human orthologues, and obtained the enrichment score (ES) based on gene set variation analysis (GSVA) as previously described^45^. Patients with breast cancer (Figure 6A), melanoma (Figure 6B), sarcoma (Figure 6C) from The Cancer Genome Atlas (TCGA) databases with a higher ES score of the LAIT specifically upregulated gene set in NK cells exhibited prolonged overall survival compared to the patients with lower ES score of such gene set expression. This result was also consistent when using upregulated genes from LAIT versus the other 3 groups combined (LAIT vs CTRL_PTT_GC) for NK cells (Figures 6D-F), indicating that LAIT upregulated gene sets may provide beneficial effect for breast cancer patient prognosis and survival extension.

**Figure 6.**
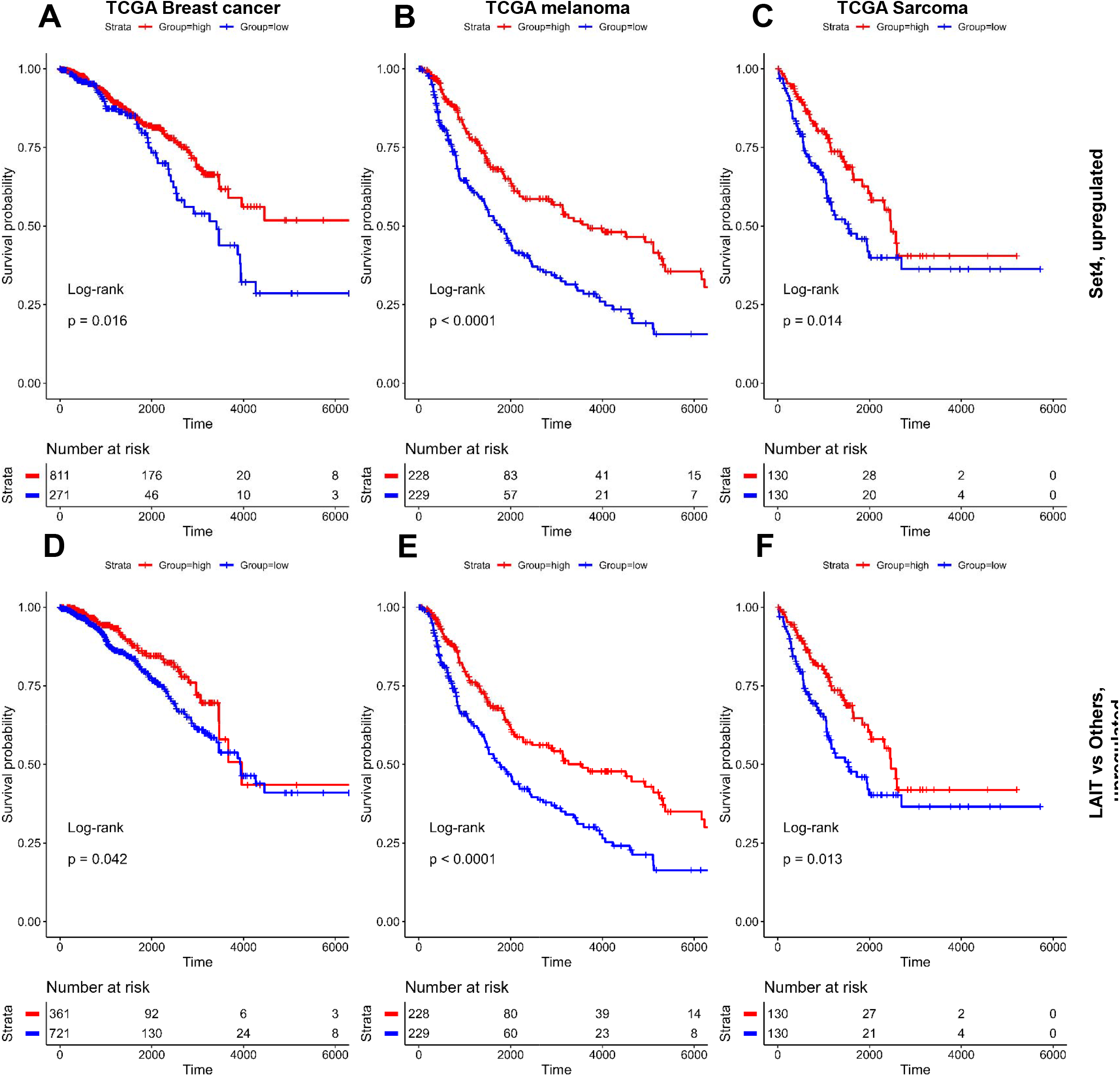
Analysis of the potential association of LAIT specifically upregulated genes with cancer patient survival. **(A)** Kaplan-Meier plots showing the significant difference in survival time (days) between breast cancer patients in groups with “high” and “low” expressions of LAIT specifically upregulated genes (Set_4 in Figure 3A). Patient groups were stratified by the first quantile of enrichment score calculated by gene set variation analysis (GSVA). Log-rank method was used for statistical analysis. **(B)** Kaplan-Meier plots showing the significant difference in survival time (days) between melanoma cancer patients in groups with “high” and “low” expressions of LAIT specifically upregulated genes (Set_4 in Figure 3A). Patient groups were stratified by the median enrichment score calculated by gene set variation analysis (GSVA). **(C)** Kaplan-Meier plots showing the significant difference in survival time (days) between sarcoma cancer patients in groups with “high” and “low” expressions of LAIT specifically upregulated genes (Set_4 in Figure 3A). Patient groups were stratified by the median enrichment score calculated by gene set variation analysis (GSVA). **(D)** Kaplan-Meier plots showing the significant difference in survival time (days) between breast cancer patients in groups with “high” and “low” expressions of LAIT vs CTRL_PTT_GC (other three groups) derived upregulated genes. Patient groups were stratified by the 2/3 of enrichment score calculated by gene set variation analysis (GSVA). **(E)** Kaplan-Meier plots showing the significant difference in survival time (days) between melanoma cancer patients in groups with “high” and “low” expressions of LAIT vs CTRL_PTT_GC (other three groups) derived upregulated genes. Patient groups were stratified by the median enrichment score calculated by gene set variation analysis (GSVA). **(F)** Kaplan-Meier plots showing the significant difference in survival time (days) between sarcoma cancer patients in groups with “high” and “low” expressions of LAIT vs CTRL_PTT_GC (other three groups) derived upregulated genes. Patient groups were stratified by the median enrichment score calculated by gene set variation analysis (GSVA).

On the other hand, the LAIT specifically downregulated genes in NK cells did not affect patient survival (Figure S6A-C). This was also consistent in LAIT vs CTRL_PTT_GC derived downregulated genes (Figures S6D-F). Taken together, we provide predictive evidence that LAIT has a greater potential to prolong patient survival by driving NK cell activation and favorable gene signatures that correlate with greater overall survival of breast cancer patients.

## Discussion

We previously reported that the combination of locally administered PTT with GC enabled induction of systemic immunity and prolonged animal survival in several aggressive tumor models ^46–49^. Our recent publications focused on the roles of tumor-infiltrating T and B cells in response to LAIT, and revealed the pro-inflammatory gene expression profiles of T cells and activation of B cells for providing anti-tumor functions^24, 50^. To further understand the mechanism of LAIT, we applied scRNA-seq analysis in this work and explored the modulation of LAIT on TINK cells, and for the first time disclosed the activation gene signatures and pathways of NK cells induced by LAIT, highlighting the potential clinical applications of GC for cancer therapy.

Specifically, we revealed by scRNAseq analysis (Figure S1A) that the PyMT TINK cells were consisted of cycling, activated, interferon-stimulated, and cytotoxic NK cell subtypes (Figure 1A-D, S1B-C), which formed a route toward activation and cytotoxicity following pseudotime progression by trajectory analysis (Figure 1E-H, S1F). Among them, LAIT mainly increased the proportion of cluster 4 of interferon NK subtype (Figure S1E), indicating the importance of interferon related functions in NK cell anti-tumor functions. PTT, GC, and LAIT shared the cytokine/interleukin-mediated signaling (Figure 2D, S2B), NK mediated cytotoxicity (Figure S2A) and TNFα signaling via NFκb (Figure 2E-G, S2A) while GC and LAIT drove the positive regulation of defense response and response to virus (Figure 2D), inflammatory response (Figure 2F-G), cytokine and receptor interaction (Figure S2C-D), and the interferon gamma signaling (Figure S2E-F), indicating the LAIT’s pro-inflammatory stimulation on NK cells. Overlapping analysis (Figure 3A, S3A) showed that LAIT specifically upregulated genes enriched in type I and II interferon pathways (Figure 3B, S3A-C) and downregulated genes enriched in protein folding/processing (Figure 4B, S4A-B) as well as heat shock protein related cell stress (Figure S4C). This was further validated by the expressions of representative genes involved in NK cell activation, cell cytotoxicity, NK stimulatory receptors, interferon pathway components, cytokine/chemokine (Figure S3D), and immune-suppressive genes, and HSP transcripts (Figure S4D). Furthermore, assessing the module score using published gene set/module associated with NK activation, cytolytic effector, pro/anti-inflammatory, type I and II interferons confirmed the anti-tumor effect of LAIT stimulated on TINK cells (Figure 5A). By performing single-cell transcriptomics analysis using immune checkpoint inhibitor (ICI)-treated mouse and human cancer patient samples, we discovered that ICI induced NK activation and cytotoxicity across several cancer types (Figure 5B, S5A-C). Furthermore, ICI-induced gene signatures showed high expression in LAIT and LAIT specifically upregulated genes demonstrated enhanced expression in ICI (Figure 5C-D, S5D-I), demonstrating that LAIT acts similarly to ICI in promoting NK cell anti-tumor activity. We also discovered that several types of cancer patients had significantly longer overall survival time if they owned higher expression of genes in NK cells that were specifically stimulated by LAIT therapy (Figure 6A-F), highlighting the potential applications of LAIT in clinical studies.

Natural killer (NK) cells are innate lymphocytes that play a critical role in the early response to infection or malignant transformation.^51^ NK cells are at the front line of efforts to develop immunological therapies due to their vigorous antitumor activity and proinflammatory roles.^52^ Under such conditions, NK cells are activated and stimulate target cell death via the release of cytolytic granules containing perforin and granzyme.^53^ NK cells express multiple inhibitory and activating receptors to discriminate and eradicate target cells, and alert the immune system by producing immunoregulatory cytokines and regulating the functions of other immune cells. ^54, 55^ The activity of NK cells is controlled by the relative balance of signals from both activating and inhibitory receptors. ^1^ Under physiological conditions, NK cell activation is inhibited by ligands, such as MHC class I (MHC I) molecules, expressed on healthy cells that recognize and engage inhibitory receptors on NK cells. Using this principle, NK cells preferentially kill MHC I-deficient tumor cells. ^56, 57^ In parallel, loss of MHC I expression has appeared as a vital mechanism by which tumor cells develop resistance to antitumor CD8^+^ T cell responses, especially in the context of immunotherapy using immune checkpoint inhibitor (ICI).^58, 59^ This indicates that the therapeutic application of NK cells may be a complemental and combinational tool for therapies that activate antitumor T cells.^60^

Immune cells within the tumor microenvironment (TME) demonstrate high heterogeneity, which is known to be modulated by many factors, including the tumor cells, stromal cells, immune cells, and endothelial cells.^61–64^ So far, flow cytometry and mass spectrometry are the main tools to characterize the heterogeneity of immune subsets and are restricted largely to cell surface molecules. ScRNA-seq is a potent technique that allows an unbiased approach without restriction to cell surface molecules, and helps in the identification of cellular landscape and diverse transcriptional signatures with single-cell resolution.^65, 66^ Recent studies using scRNAseq identified the landscape of NK cell subtypes and gene signatures from both healthy and tumor tissues in mouse and human.^67–72^ Our work provides a transcriptional atlas of individual NK cells from mouse breast tumor model, revealing distinct features of tumor-infiltrating NK cells upon LAIT treatment. Consistent with these scRNAseq-based findings for NK cell subsets, we identified the *Cd27^+^Itgam^-^*and *Cd27^-^Itgam^+^* (cluster 2) NK cells. We also found that the *Cd27^-^ Itgam^+^* (cluster 2) NK cells displayed high expression of *Gzma/b*, *Prf1*, *Ncr1*, *Klrk1*, and *Ifng* and low expression of *Lag3*. Our finding of activated NK cell signatures was also in agreement with a recent report from activated Hif1a^−/−^ NK cells^73^, demonstrating the capacity of scRNAseq in characterizing the NK cell subset transcriptomics.

Numerous studies have found that TINK cells are dysfunctional due to the immunosuppressive microenvironment^1, 74^. Therefore, TINK cells display impaired anti-tumor phenotype and poor cytotoxic function in many tumors, including breast cancer^75^, melanoma^76^, lung cancer,^77^ and liver cancer.^78^ The conversion of effector NK cells into type 1 ILCs mediated by immunosuppressive TGF-β has also been demonstrated as a mechanism of tumor evasion from NK cell cytotoxicity in the TME.^79, 80^ We found LAIT significantly inhibited *Tgfb* related signaling and therefore contributes to release of such immunosuppression.

Accumulating evidences highlighted the acquisition of immune checkpoint molecules by TINK cells associate with the achievement of a regulatory phenotype, characterized by T cell suppression mediated by expression of immune checkpoint molecules. For example, Expression of CTLA4 ^81^, PD-1^82^, LAG3^83^, TIGIT^84^ in NK was found to be associated with NK cell exhaustion in multiple cancer types in both tumor-bearing mice and patients. In our work, the finding that only CTLA4 and LAG3, while not other checkpoints were expressed in NK cells, confirmed the tumor suppressive function of NK cells in the PyMT TME. More importantly, LAIT could decrease the expression of LAG3, implying that it can somewhat counteract immune suppression through such checkpoint. We also note that CTLA4 could not be inhibited while rather promoted, which was similarly found in PyMT T cells^50^. This reminds us to adopt combination of LAIT with anti-CTLA4 for a potentially achieving better cancer therapeutic efficacy.

NK cell numbers have been shown to be correlated with survival of patient with multiple cancer types. For example, CD56^+^ NK cells were found associated with a better outcome in head and neck squamous cell carcinoma^85^, renal cell carcinoma^86^ and colorectal carcinoma^87^. In addition to NK cell number, the NK cell activity seems to be more helpful to determine the real influence of NK cells on clinical outcome^87^. The importance of NK cell activity was highlighted by a study that showed a correlation between the low expression of NKp30/Ncr3 genes and a poor clinical outcome.^88^ Another similar work analyzed the cytotoxic activity of peripheral NK cells among 154 cancer cases and found an inverse correlation between NK cell cytotoxic activity and cancer incidence.^89^ In our work, we also found treatment especially LAIT strongly upregulated the expressions of a panel of activation/effector molecules, type I/II interferon genes, and activating receptors including above mentioned *Ncr1*. This NK activation-supporting proinflammatory expression profiles laid the foundation for the positive correlation with favorable prognosis and overall survival in several cancer types in our TCGA data analysis.

The theranostic value of NK cells has recently been investigated with respect to immune checkpoints inhibitor (ICI) therapy, as immunotherapies has become the most recent and remarkable improvement of cancer treatment. In melanoma, lymphoma and colon carcinoma murine models, it was demonstrated that the PD-1/PD-L1 axis enabled modulation of NK cell phenotype, as such blockade restored NK cell antitumor functions and achieved a better survival of the animals.^90^ The use of an anti-PD-1 antibody Nivolumab in lung cancer patients increased the NK cell number in patients with better prognosis.^91^ Similarly, NCR1 was the parameter affecting the prognosis in non-small cell lung carcinoma (NSCLC) patients with high PD-L1 expression on tumor cells.^92^ In metastatic melanoma, the increased density of activated NK cells into the tumor was correlated with response to PD-1 blockade, concurrent with activation of NK cell cytotoxicity.^93^ NK cell activation was also found predictive to non-progressive disease and high progression free survival (PFS) in patients with melanoma, lung cancer and head and neck cancers, in response to anti-PD-1 therapy.^94^ Our scRNAseq analysis using human cancer patients receiving ICIs showed that such blockade induced an activation and effector proinflammatory phenotype and enriched type I/II pathways in a series of cancer types. We also found LAIT and ICIs achieved similar effect in activating NK cell transcriptomics. This comparative analysis for the first time bridges the similarity of LAIT and ICI and the mechanism will be studied in the future work.

Overall, this study provided novel insights into how LAIT-induced NK cell activation and effector function and its specifically upregulated genes showed positive correlation with prolonged cancer patient survival. These findings revealed the similarities of LAIT and ICI-based immunotherapies in NK cell antitumor activity enhancement, broadened our understanding of LAIT’s modulatory roles in remodeling TME NK cells and shed light on the potential of NK cell activation in clinical applications.

## Abbreviations

CTRL: control
PTT: photothermal therapy
GC: N-dihydrogalactochitosan
LAIT: localized ablative immunotherapy
MMTV-PyMT: mouse mammary tumor virus-polyoma middle tumor-antigen
scRNA-seq: single-cell RNA sequencing
TILs: tumor-infiltrating lymphocytes
TINKs: tumor-infiltrating NK cells
TME: tumor microenvironment
IFN-γ: interferon-γ
TNFα: tumor necrosis factor-α
t-SNE: t-distributed stochastic neighbor embedding
UMAP: uniform manifold approximation and projection
DEGs: differentially expressed genes
ES: enrichment score
GO: gene ontology
ORA: over-representation analysis
GSEA: gene set enrichment analysis
GSVA: gene set variation analysis
KEGG: kyoto encyclopedia of genes and genomes
MsigDB: Molecular Signatures Database.

## Methods

### Sample preparation and data processing for single-cell RNA sequencing

Tumor sample processing, tumor-infiltrating immune cell preparation, and single-cell RNA sequencing barcoding and library generation were performed according to our previous work.^24, 50^ FACS sorted live CD45^+^ tumor-infiltrating immune cells were encapsulated into droplets via a 10× Genomics platform according to the manufacturer’s instructions. Paired-end RNA-seq was performed via an Illumina NovaSeq 6000 sequencing system. The Linux command *cellranger count* from *Cell Ranger* pipeline was used to process sample-specific *FASTQ* files into feature-barcode matrices. *Seurat* (v3.2) was used to process and integrate above scRNAseq datasets from different treatment groups.^24, 32, 95, 96^ Total CD45^+^ immune cells were quality controlled, normalized, integrated, clustered and lymphoid cell clusters were obtained according to our previous work. Then, T cells with expressions of *Cd3d*, *Cd3g*, *Cd3e*, *Cd8a*, *Cd8b1*, *Cd4*, *Tcrg- C1*, and B cells with expression of *Cd19*, were removed. Then, Natural killer (NK) cells were remained with expressions of *Ncr1* (CD335/NKp46) or Klrb1c (CD161/NK1.1). These NK cells were re-clustered with resolution 0.3 to generate its UMAP plot.

### Single-cell trajectory analysis

*Monocle2* R package was chosen for single-cell trajectory inference using *DDRTree* algorithm.^26^ The tumor-infiltrating CD8^+^ and CD4^+^ T cell *Seurat* objects were imported into *Monocle2* and highly variable genes calculated by *differentialGeneTest* function were used for gene ordering and trajectory construction.

### Differential gene expression (DGE) and functional enrichment analysis

Treatments of PTT, GC, and LAIT (PTT+GC) were compared with CTRL, respectively. *FindMarkers* function in *Seurat* R package was used to obtain differential gene expression for each comparison with threshold of Log fold change (logFC) set at 0.25 and −0.25 as default.^32^ Top 10 of both upregulated and downregulated differentially expressed genes (DEGs) were labelled on volcano plots visualized by using *ggplot2* in R. *clusterProfiler* R package was adopted for gene functional enrichment analysis by using both over-representation analysis (ORA) and gene set enrichment analysis (GSEA) methods and gene annotation and pathway databases including GO, KEGG, Reactome and MsigDB.^35^ Dot plot and network plot were mainly used for ORA and GSEA result visualizations respectively. GSEA obtained normalized enrichment scores (NES) for pathways were used for clustering in heatmap.

### Overlapping of treatment-derived DEGs

Treatment-derived DEGs were overlapped and visualized by using *VennDiagram* R package following with subset of indicated gene sets functional enrichment analysis.^35^ Expression of these subset genes were obtained using *AverageExpression* function in *Seurat* R package^32^ and visualized using circular heatmap.^97, 98^

### ScRNAseq analysis for NK cell transcriptomics from immune checkpoint inhibitor treated cancer patients and mouse

ScRNAseq data of the tumor and blood samples from melanoma patients of CY129, CY155, CY158, CY160, CY164 were used for study the NK cell atlas.^41^ CY160 was the patient without ICI treatment while other four patients were upon ICI therapy according to the author’s descriptions. Although the author claims all these cells were NK cells, we conducted data cleaning by excluding potential non-NK cells by using the markers of T cell (CD3G, CD3D, CD3E), B cell (CD19, MS4A1), Myeloid cell (CD14, FUT4, CD163) and obtained 22818 NK cells.

To study the effect of ICI on NK cells in a mouse breast cancer model, the scRNAseq dataset from breast cancer mice upon ICI (anti-PD1/anti-CTLA4) treatments for 7 days were used.^42^ NK cells from both T11-Apobec model and KPB25L-UV model were subset and filtered similarly to the descriptions of PyMT NK cells in our work. After obtaining 94 NK cells from KPB25Luv model and 662 NK cells from T11-Apobec model, the latter was kept for downstream analysis.

The scRNAseq data from breast cancer patients upon ICI (anti-PD1) treatment were also analyzed.^43^ According to the author’s annotated metadata, the “Pre” labeled patients refers to non-ICI treatments and “On” refers to ICI treatment. Both cytotoxic NK cells (NK_CYTO) and resting NK cells (NK_REST) were selected as NK cells. 1298 NK cells from patients with clonally expanded T cells were used for downstream analysis.

The scRNAseq dataset from leptomeningeal metastases patients upon ICI (anti-PD1 antibody pembrolizumab, anti-CTLA4 antibody ipilimumab) treatments were analyzed for the purpose of investigating the effect of ICI on NK cells.^44^ According to the author’s annotated metadata, the “pre” labeled patients refers to non-ICI treatments and “post” refers to ICI treatment. NK cells were selected similar to the criteria we used for breast cancer patients. 1432 NK cells from patients were used for downstream analysis.

### Correlation between clinical survival and expression of given gene set

Gene expression RNAseq and survival data available from GDC TCGA Breast Cancer (BRCA), melanoma, and sarcoma were downloaded from UCSC Xena (https://xenabrowser.net/datapages/) and used for assessing the clinical significance of expression of a given gene set.^99^ To evaluate whether there was a relationship between differentially expressed gene set and clinical survival in breast cancer patients, we utilized both LAIT specifical DEGs and LAIT vs CTRL_PTT_GC combined groups to obtain the upregulated and downregulated gene sets and visualized in volcano plot. Gene symbols in mouse were converted to gene symbols in human by using *biomaRt* R package.^100^ These gene sets were taken as the input to get the enrichment score (ES) using gene set variation analysis (GSVA) R package.^45^ Cancer patients were stratified as the “high” group if ES was above the indicated cutoff (such as median or ¼) and as “low” group if ES is below the cutoff. Analyses were performed with Kaplan-Meier estimates and log-rank tests. Numbers below plots represent numbers of breast cancer patients and survival time are days.

### Statistical analyses

Statistical analysis was conducted with R. One-way analysis of variance (ANOVA) was used for multiple group comparisons. Proportion test function (“prop.test”) in R was used for cell proportion test. Kruskal-Wallis test for multiple comparison and Wilcoxon test for two group comparison were provided by the R package *ggpubr*. Differences in survival were determined on the basis of Kaplan-Meier survival analysis. Adjusted *P* values less than or equal to 0.05 were considered statistically significant (*, *p* ≤ 0.05; **, *p* ≤ 0.01).

## Data and code availability

The accession number for the scRNAseq data reported in this paper is GEO: GSE150675.^24, 50^ Analysis for such data can be available upon request.

## Acknowledgments

This work was supported in part by the National Cancer Institute (R01CA205348 and R01CA269897 to WRC), National Institute of General Medical Sciences (P20GM135009), and the Oklahoma Center for the Advancement of Science and Technology (HR16-085 and HF20-019 to WRC). We would like to thank the Flow Cytometry Core Facility and Genomics Core Facility at the Oklahoma Medical Research Foundation (OMRF) for providing flow cytometry services and assisting in scRNAseq experiment.

## Ethics declarations

Wei R. Chen is co-founder and an unpaid member of the Board of Directors of Immunophotonics, Inc. All other authors declare no competing interests associated with this publication.

## Notes

### Competing Interest Statement

The authors have declared no competing interest.

