## Supplementary material for "Single-cell transcriptomics reveals that tumor-infiltrating natural killer cells are activated by localized ablative immunotherapy and share anti-tumor signatures induced by immune checkpoint inhibitors": suppl figure legends

**Supplemental Figure Legends**

**Figure S1.** **Schematic** **of treatment and scRNAseq analysis for tumor-infiltrating NK cells (related to Figure 1).**

**(A)** Schematic of MMTV-PyMT tumor implantation, animal treatment, sample collection, and scRNAseq preparation and bioinformatics analysis. DGE: differential gene expression.

**(B)** Feature plots showing specific expressions of selected NK cell markers.

**(C)** Heatmap showing expressions for the top 10 highly expressed cluster marker genes.

**(D)** UMAP plots of NK cells in different treatment groups (CTRL, PTT, GC, and LAIT).

**(E)** Bar plot showing the NK cell proportion in each cluster in different treatment groups (CTRL, PTT, GC, and LAIT). The proportion was calculated by using NK cell number in each cluster divided by the total CD45^+^ immune cells within given treatment.

**(F)** Heatmap showing expressions of selected NK cell marker genes that significantly change following pseudotime progression.

**Figure S2.** **Differential gene expression and pathway enrichment analysis for TINK cells (related to** **Figure 2)**.

**(A)** Dot plot showing KEGG enrichment analysis using ORA method for both upregulated and downregulated genes from comparisons of PTT vs CTRL, GC vs CTRL and LAIT vs CTRL.

**(B)** Dot plot showing Reactome enrichment analysis using ORA method for both upregulated and downregulated genes from comparisons of PTT vs CTRL, GC vs CTRL and LAIT vs CTRL.

**(C)** Network plot of the KEGG enrichment analysis using GSEA method for DEGs from GC vs CTRL.

**(D)** Network plot of the KEGG enrichment analysis using GSEA method for DEGs from LAIT vs CTRL.

**(E)** Network plot of the Reactome enrichment analysis using GSEA method for DEGs from GC vs CTRL.

**(F)** Network plot of the Reactome enrichment analysis using GSEA method for DEGs from LAIT vs CTRL.

**Figure S3. Overlapping of treatment-upregulated genes in TINK cells (related to** **Figure 3).**

**(A)** Dot plot for MsigDB enrichment analysis of genes in Set_1 to Set_4.

**(B)** Dot plot for KEGG enrichment analysis of genes in Set_1 to Set_4.

**(C)** Dot plot for Reactome enrichment analysis of genes in Set_1 to Set_4.

**(D)** Heatmap showing the expression of selected genes in each treatment group in each NK cell subtype.

**Figure S4. Overlapping of treatment-downregulated genes in TINK cells (related to** **Figure 4).**

**(A)** Dot plot for MsigDB enrichment analysis of genes in Set_1 to Set_4.

**(B)** Dot plot for KEGG enrichment analysis of genes in Set_1 to Set_4.

**(C)** Dot plot for Reactome enrichment analysis of genes in Set_1 to Set_4.

**(D)** Heatmap showing the expression of selected immune suppressive genes in each treatment group in each NK cell subtype.

**Figure S5.** **ScRNAseq analysis for similarity between LAIT and immune checkpoint inhibitor (ICI) in stimulating anti-tumor phenotype of NK cells (related to Figure 5).**

**(A)** Violin plot showing module scores for anti-tumor phenotype related gene signatures including NK activation, cytolytic effector, pro-inflammatory, anti-inflammatory, type I interferon, and type II interferon gene signatures in ICI treated TINK cells from a high mutation burden mouse model of breast cancer (Hollern et al., 2019; GSE136206).

**(B)** Violin plot showing module scores for anti-tumor phenotype related gene signatures including NK activation, cytolytic effector, pro-inflammatory, anti-inflammatory, type I interferon, and type II interferon gene signatures in ICI treated TINK cells from breast cancer patients (Bassez et al., 2021; a download of the read count data per individual patient is publicly available at <http://biokey.lambrechtslab.org>). The labels of ‘Pre’ and ‘On’ were from the author’s annotated data. ‘Pre’ represents patient samples before ICI treatment; ‘On’ represents patient samples with/upon ICI treatment.

**(C)** Violin plot showing module scores for anti-tumor phenotype related gene signatures including NK activation, cytolytic effector, pro-inflammatory, anti-inflammatory, type I interferon, and type II interferon gene signatures in ICI treated TINK cells from patients with leptomeningeal metastases disease (Prakadan et al., 2021; Processed scRNA data were downloaded from single cell portal with study number SCP1332). The labels of ‘pre’ and ‘post’ were from the author’s annotated data. ‘pre’ represents patient samples before ICI treatment; ‘post’ represents patient samples with/upon ICI treatment.

**(D)** Violin plot showing module scores for ICI-induced gene signatures mapped into treatment groups of CTRL, PTT, GC, and LAIT in mouse PyMT TINK cells. ICI-induced gene signatures from mouse breast cancer model T11.Apobec NK cells demonstrated the increased production in GC/LAIT group, indicating the similar effect of ICI and LAIT.

**(E)** Violin plot showing module scores for LAIT specifically-induced gene signatures (upregulated Set_4) mapped into mouse breast cancer model T11.Apobec NK cells upon ICI treatment. LAIT specifically-induced gene signatures from PyMT mouse TINK cells demonstrated the increased production in ICI treatment group, further confirming the similar effect of ICI and LAIT in stimulating NK cell’s anti-tumor immunity.

**(F)** Violin plot showing module scores for ICI-induced gene signatures mapped into treatment groups of CTRL, PTT, GC, and LAIT in mouse PyMT TINK cells. ICI-induced gene signatures from breast cancer patient TINK cells demonstrated the increased production in LAIT group when compared with untreated CTRL, indicating the similar effect of ICI and LAIT.

**(G)** Violin plot showing module scores for LAIT specifically-induced gene signatures (upregulated Set_4) mapped into breast cancer patient TINK cells upon ICI treatment. LAIT specifically-induced gene signatures from PyMT mouse TINK cells demonstrated the increased production in ICI treatment group, further confirming the similar effect of ICI and LAIT in stimulating NK cell’s anti-tumor immunity.

**(H)** Violin plot showing module scores for ICI-induced gene signatures mapped into treatment groups of CTRL, PTT, GC, and LAIT in mouse PyMT TINK cells. ICI-induced gene signatures from mouse breast cancer model T11.Apobec NK cells demonstrated the increased production in GC/LAIT group, indicating the similar effect of ICI and LAIT.

**(I)** Violin plot showing module scores for LAIT specifically-induced gene signatures (upregulated Set_4) mapped into patients with leptomeningeal metastases disease. TINK cells upon ICI treatment. LAIT specifically-induced gene signatures from PyMT mouse TINK cells demonstrated the increased production in ICI treatment group, further confirming the similar effect of ICI and LAIT in stimulating NK cell’s anti-tumor immunity.

**Figure S6. Analysis of the potential association of LAIT specifically upregulated genes with cancer patient survival (related to Figure 6).**

**(A)** Kaplan-Meier plots showing the insignificant difference in survival time (days) between breast cancer patients in groups with “high” and “low” expressions of LAIT specifically downregulated genes (Set_4 in Figure 4A). Patient groups were stratified by the first quantile of enrichment score calculated by gene set variation analysis (GSVA). Log-rank method was used for statistical analysis.

**(B)** Kaplan-Meier plots showing the insignificant difference in survival time (days) between melanoma cancer patients in groups with “high” and “low” expressions of LAIT specifically downregulated genes (Set_4 in Figure 4A). Patient groups were stratified by the median enrichment score calculated by gene set variation analysis (GSVA).

**(C)** Kaplan-Meier plots showing the insignificant difference in survival time (days) between sarcoma cancer patients in groups with “high” and “low” expressions of LAIT specifically downregulated genes (Set_4 in Figure 4A). Patient groups were stratified by the median enrichment score calculated by gene set variation analysis (GSVA).

**(D)** Kaplan-Meier plots showing the insignificant difference in survival time (days) between breast cancer patients in groups with “high” and “low” expressions of LAIT vs CTRL_PTT_GC (other three groups) derived downregulated genes. Patient groups were stratified by the 2/3 of enrichment score calculated by gene set variation analysis (GSVA).

**(E)** Kaplan-Meier plots showing the insignificant difference in survival time (days) between melanoma cancer patients in groups with “high” and “low” expressions of LAIT vs CTRL_PTT_GC (other three groups) derived downregulated genes. Patient groups were stratified by the median enrichment score calculated by gene set variation analysis (GSVA).

**(F)** Kaplan-Meier plots showing the insignificant difference in survival time (days) between sarcoma cancer patients in groups with “high” and “low” expressions of LAIT vs CTRL_PTT_GC (other three groups) derived downregulated genes. Patient groups were stratified by the median enrichment score calculated by gene set variation analysis (GSVA).
