## Supplementary material for "Single-cell transcriptomics reveals that tumor-infiltrating natural killer cells are activated by localized ablative immunotherapy and share anti-tumor signatures induced by immune checkpoint inhibitors": suppl figures

### Slide 1
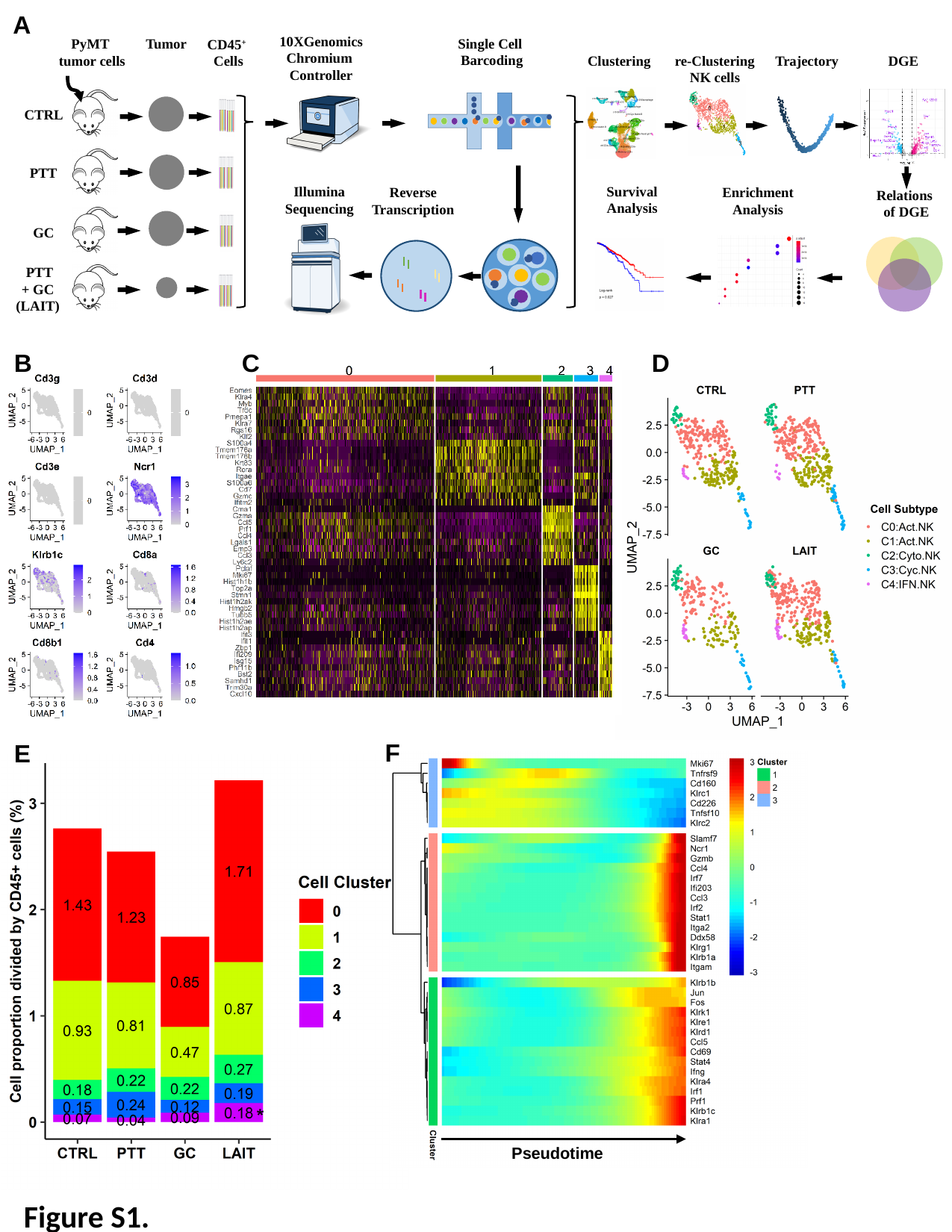

A
10XGenomics
Chromium
Controller
Single Cell
Barcoding
PyMT
tumor cells
Tumor
CD45^+^
Cells
Clustering
re-Clustering
NK cells
Trajectory
DGE
CTRL
PTT
Survival
Analysis
Illumina
Sequencing
Enrichment
Analysis
Reverse
Transcription
Relations
of DGE
GC
 PTT
+ GC
(LAIT)
C
D
B
F
E
Pseudotime
Figure S1.

### Slide 2
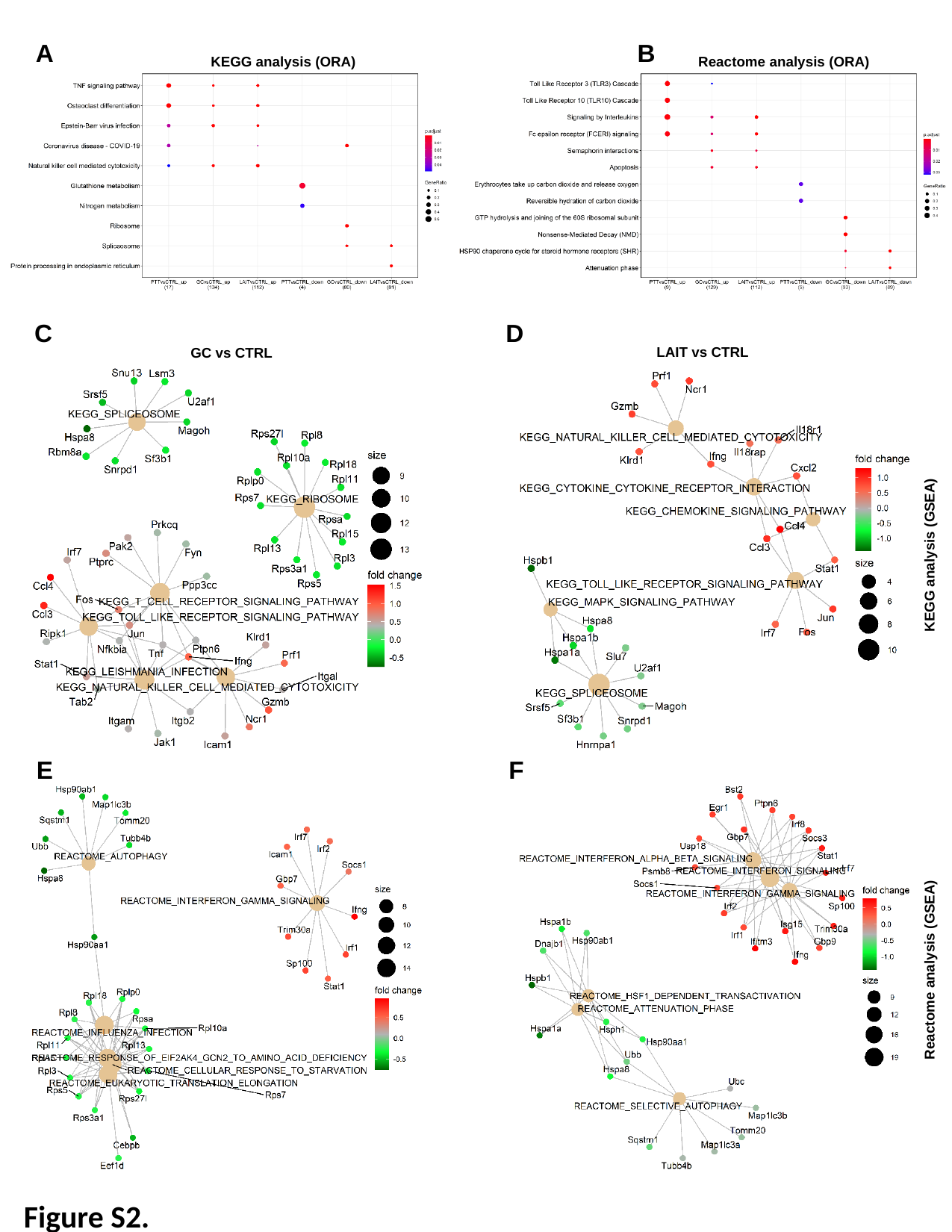

A
B
KEGG analysis (ORA)
Reactome analysis (ORA)
C
D
LAIT vs CTRL
GC vs CTRL
KEGG analysis (GSEA)
E
F
Reactome analysis (GSEA)
Figure S2.

### Slide 3
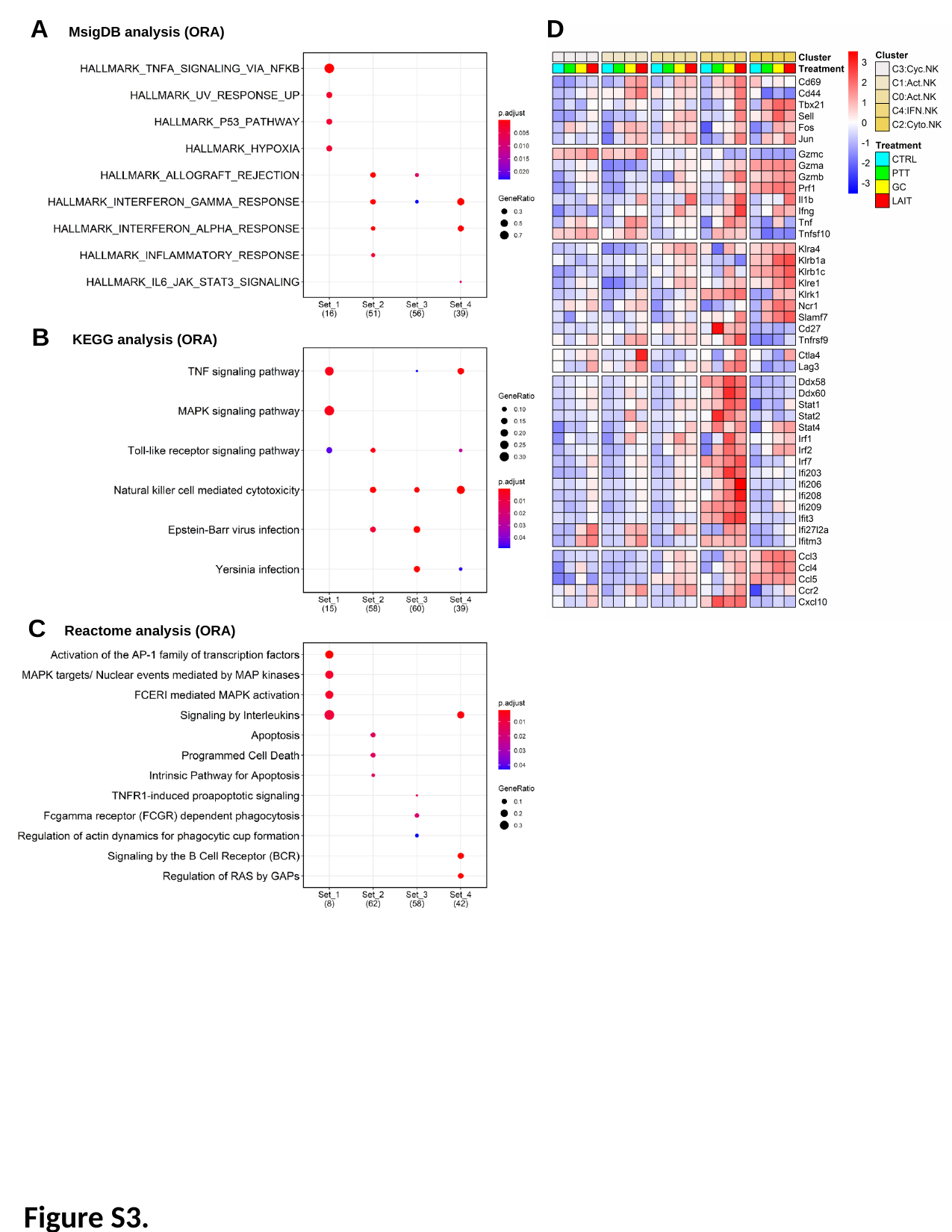

A
D
MsigDB analysis (ORA)
B
KEGG analysis (ORA)
C
Reactome analysis (ORA)
Figure S3.

### Slide 4
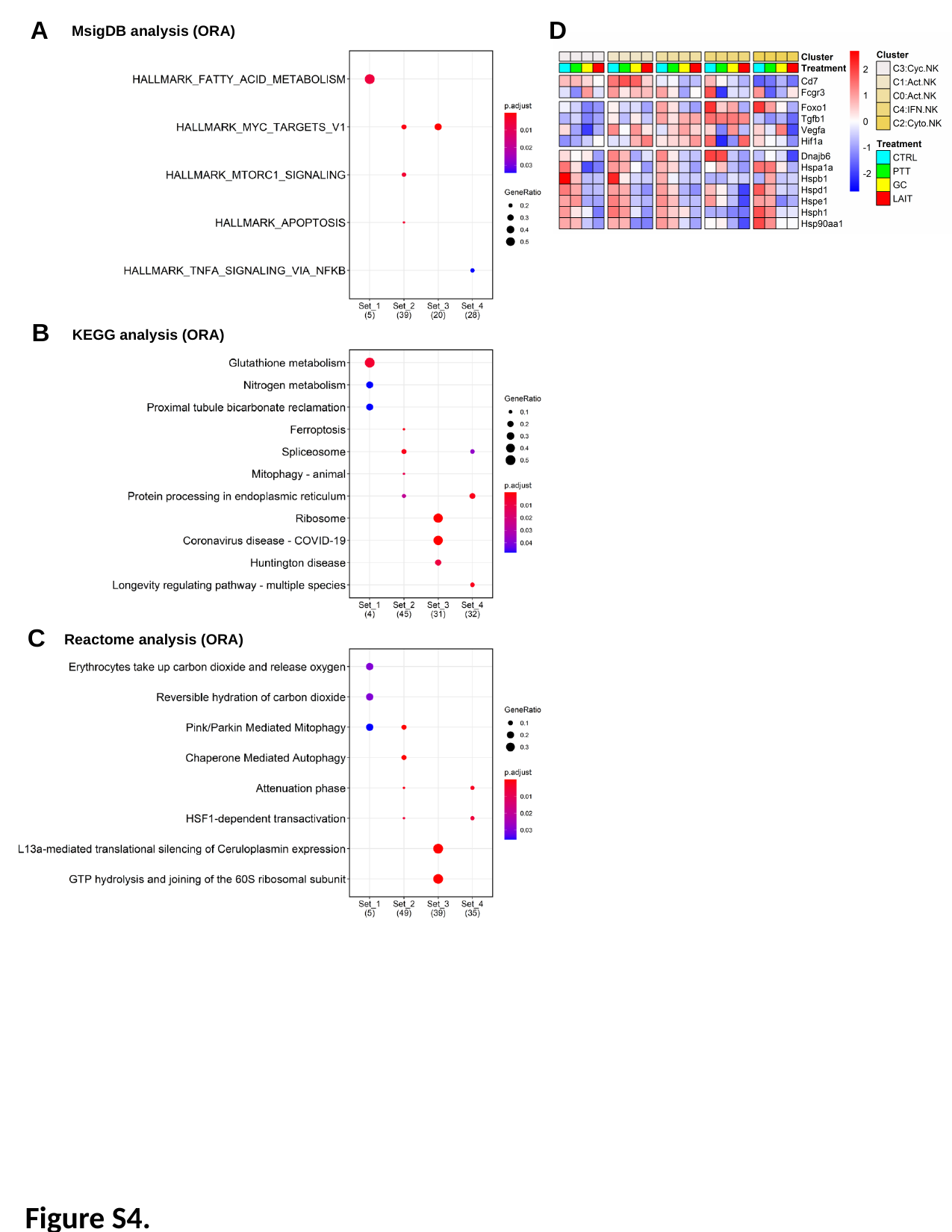

A
D
MsigDB analysis (ORA)
B
KEGG analysis (ORA)
C
Reactome analysis (ORA)
Figure S4.

### Slide 5
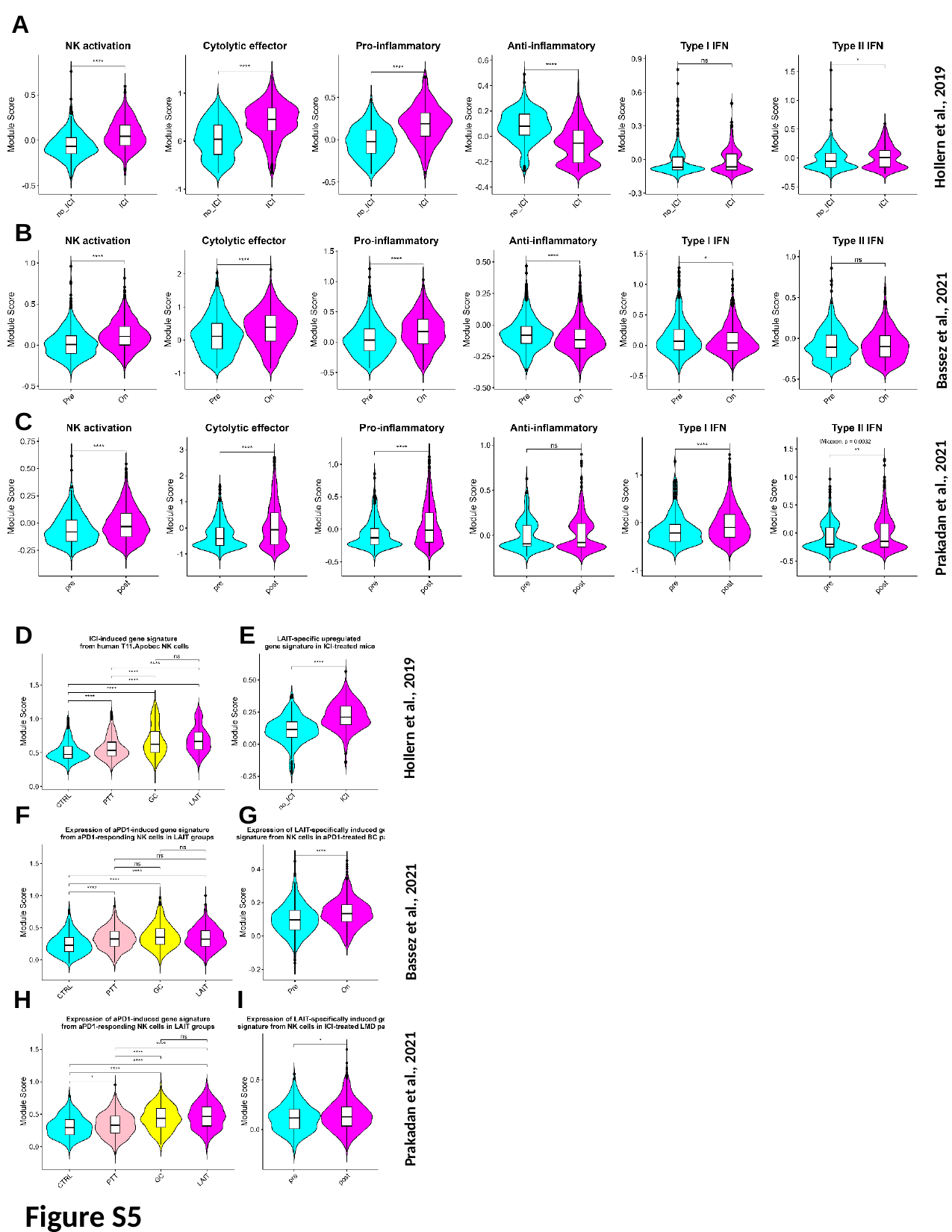

A
GSE136206
Hollern et al., 2019
B
Bassez et al., 2021
C
Prakadan et al., 2021
SCP1332
D
E
Hollern et al., 2019
F
G
Bassez et al., 2021
H
I
Prakadan et al., 2021
Figure S5

### Slide 6
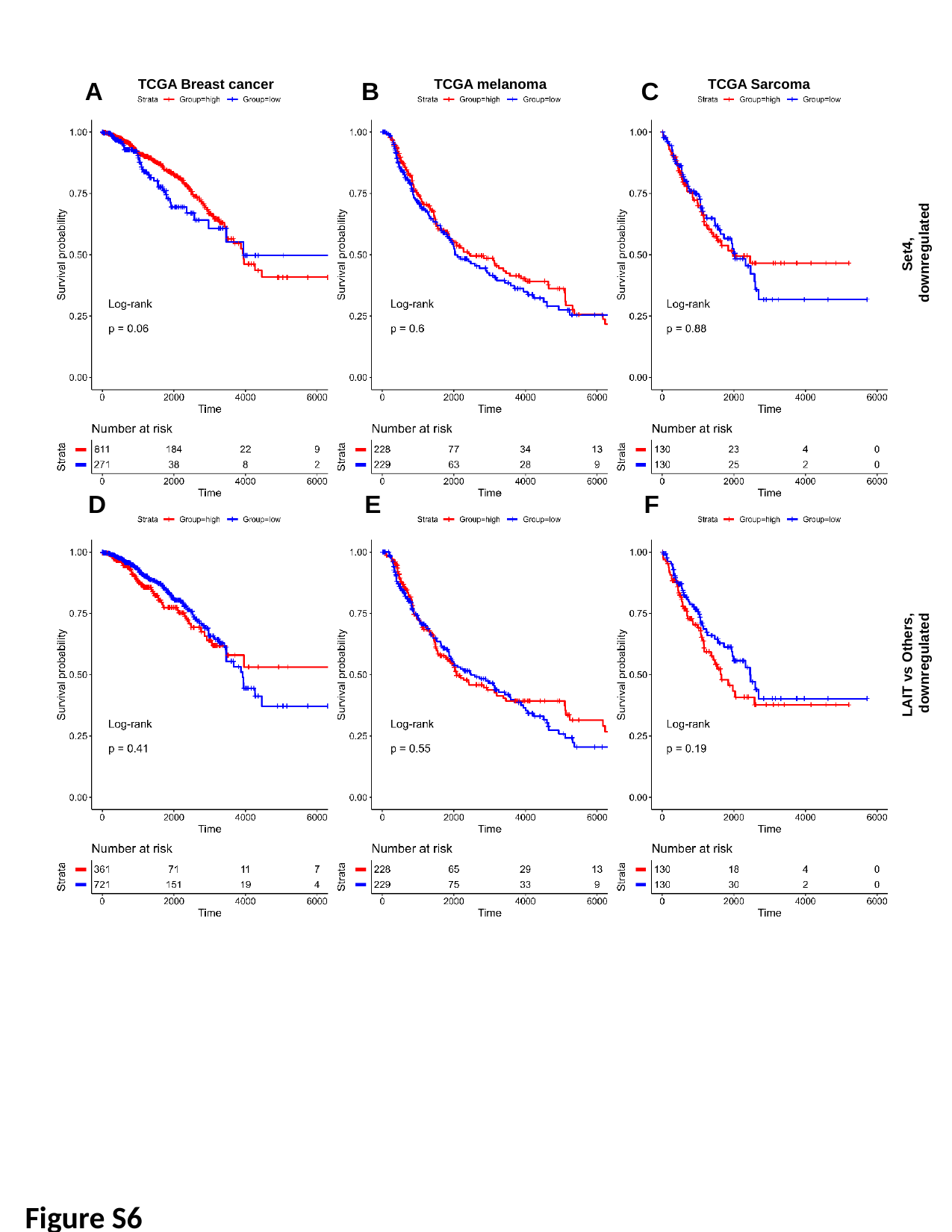

A
TCGA Breast cancer
TCGA melanoma
C
TCGA Sarcoma
B
Set4,
downregulated
D
F
E
LAIT vs Others,
downregulated
Figure S6
